# Global anti-tumor immunity after localized, bioengineered Treg depletion

**DOI:** 10.1101/2021.09.02.458797

**Authors:** Fatemeh S. Majedi, Mohammad Mahdi Hasani-Sadrabadi, Timothy J. Thauland, Sundeep G. Keswani, Song Li, Louis-S. Bouchard, Manish J. Butte

**Author notes:** Corresponding Author: Manish Butte, MD PhD, 10833 Le Conte Ave, MDCC Building Room 12-430, Los Angeles, CA 90095.

## Abstract

Over 90% of deaths from cancer occur due to solid tumors, occurring at a rate of ∼1,500 deaths per day in the US, highlighting a profound and unmet need for new therapies. Solid tumors evade clearance by T cells due to a variety of immunosuppressive properties of the tumor microenvironment. However, this immunosuppression cannot be easily blocked on a global level because systemic activation of the immune system elicits a host of complications. An ideal therapy for solid tumors would act *locally* to activate the immune response without evoking global adverse effects. Here we present a biodegradable, macroporous scaffold that is implanted adjacent to the tumor and suppresses the main obstacle to cancer immunosurveillance: intratumoral regulatory T cells. The scaffold also promotes the recruitment and activation of T cell effectors into the tumor, resulting in clearance of otherwise aggressive and fatal tumors in mice. Unexpectedly, the local depletion of Tregs results in an “immunological abscopal effect” acting on distant tumors. We demonstrate that this versatile platform can also deliver tumor-antigen-specific T cells directly to the peri-tumoral environment, bypassing difficulties in intravenous delivery including the environmental barriers imposed by the tumor’s vasculature. By orchestrating multiple local immunomodulatory treatments, this scaffold offers a general approach to engineer T-cell responses to solid tumors without systemic toxicities.

---

Solid tumors effectively evade immune surveillance by modifying the local environment in such a way that the immune response is suppressed. Systemic solutions to overcoming the immunosuppressive tumor microenvironment have serious downsides: for example, global inhibition of regulatory T cells can lead to autoimmune side effects whereas systemic activation of T cells can lead to cytokine storms. Thus, no good solutions currently exist to the problem of global regulation of the immune response targeting tumors. Immunomodulatory solutions that act locally, thereby avoiding global side effects, would be attractive. Here we developed a biodegradable scaffold (sponge-like biomaterial) that can be placed locally adjacent to the tumor bed to robustly activate endogenous T-cell immunity against the tumor. Our scaffold comprises many design considerations that enhance local immunity against tumors. First, it blocks a key growth factor for regulatory T cells. Second, it entices effector T cells into the microenvironment and into the microporous structure of our scaffold using chemokines. Third, it super-activates effector T cells through stimulatory antibodies. Fourth, we tuned the mechanical stiffness of the scaffold to maximize mechanosensing pathways in T cells. Fifth, the scaffold secretes stimulatory cytokines to promote potent T-cell activation and the development of memory T cells. All five strategies are needed to obtain the highly effective antitumor response presented herein.

TGF-β is a potent growth factor of the tumor microenvironment that promotes cancer proliferation, metastasis, and the induction of regulatory T cells (Treg) from the helper T cells drawn to the tumor. TGF-β also inhibits the effector functions and the migration of cytotoxic T cells in the tumor.^1,2^ TGF-β has thus become a popular target for immunotherapy.^3^ However, systemic TGF-β inhibition in preclinical models and humans has shown major adverse effects on the cardiovascular, gastrointestinal, and skeletal systems.^4,5^ The release of TGF-β inhibitors by injected nanoliposomes reduced metastases,^6^ but did not reduce regulatory T cells locally, a desirable condition for fighting solid tumors. Thus, local suppression of regulatory T cells appears necessary but remains elusive.

Promoting the effector functions of endogenous, tumor-associated T cells is a major goal of immunotherapies. The release of cytokines to T cells via nanogel “backpacks” was shown to enhance tumor clearance.^7^ However, the continual and uncontrolled release of cytokines risks systemic effects. Intravenous administration of collagen-binding domains fused to the cytokine IL-12 prolonged the release of cytokine into the tumor stroma^8^; however, this approach increases risks of cardiovascular disease. We have previously developed nanoparticles that deliver cytokines to CD8+ T cells at carefully controlled rates and showed that prolonged exposure to IL-2 impacts their differentiation and effector function.^9^ Besides cytokines, antibodies that stimulate CD3 and CD28 are known to augment T cell activation *in situ*. Intratumoral injection of stimulatory antibodies has been attempted but rapid transport away from the tumor environment has prevented potential clinical application.^10,11^ Our approach is to anchor stimulatory antibodies and prolong cytokine release in the scaffold.

In addition to biochemical cues, we showed in recent work that T cells sense the mechanical stiffness of 3D microenvironments and undergo augmented activation when the environment reaches an elastic modulus of ∼40 kPa.^12^ Mechanistically, we have shown that the stiffness of the local environment affects T-cell activation by modulating their metabolic program.^13^ Others have employed biomechanically soft hydrogels *in vivo* to activate T cells,^14^ but we now know that mechanical rigidity is important to maximize activation. The mechanical stiffness of our biomaterial is optimized to mimic that of activated lymph nodes^14^, hence its key role in our approach.

To facilitate the immune response against solid tumors, we introduce a new multifunctional biomaterial scaffold that is implanted adjacent to a tumor, which attracts cytotoxic T cells and enhances their function by suppressing tumor-resident Tregs. Together these activities allow for the much sought-after effect of overcoming the immunosuppressive effects of the microenvironment of solid tumors, a much sought-after goal of solid tumor immunotherapy. This flexible platform offers a new promise for localized immunomodulation and treatment of cancer. The methodology is general and could be adapted for localized immunoregulation in autoimmune diseases and infections as well.

## Results

### Scaffold fabrication

To enhance local immunity against tumors, our scaffold combines five design elements (**Fig. 1a**). First, the IL-2 cytokine is gradually eluted from engineered mesoporous silica microparticles (**Extended Data Fig. 1, Extended Data Fig. 3b**). Second, TGF-β inhibitor is gradually released from embedded PLGA nanoparticles (**Extended Data Fig. 2**), which suppresses Tregs (**Extended Data Fig. 3g-i**). Third, chemokines to attract T cells are gradually released from alginate scaffolds (**Extended Data Fig. 4**). Fourth, the mechanical stiffness of the microporous scaffold was tuned to maximize T-cell activation (**Extended Data Fig. 5**). Fifth, surfaces of the scaffold were modified with T-cell stimulatory antibodies (**Extended Data Fig. 3c-f**).

**Fig. 1.**
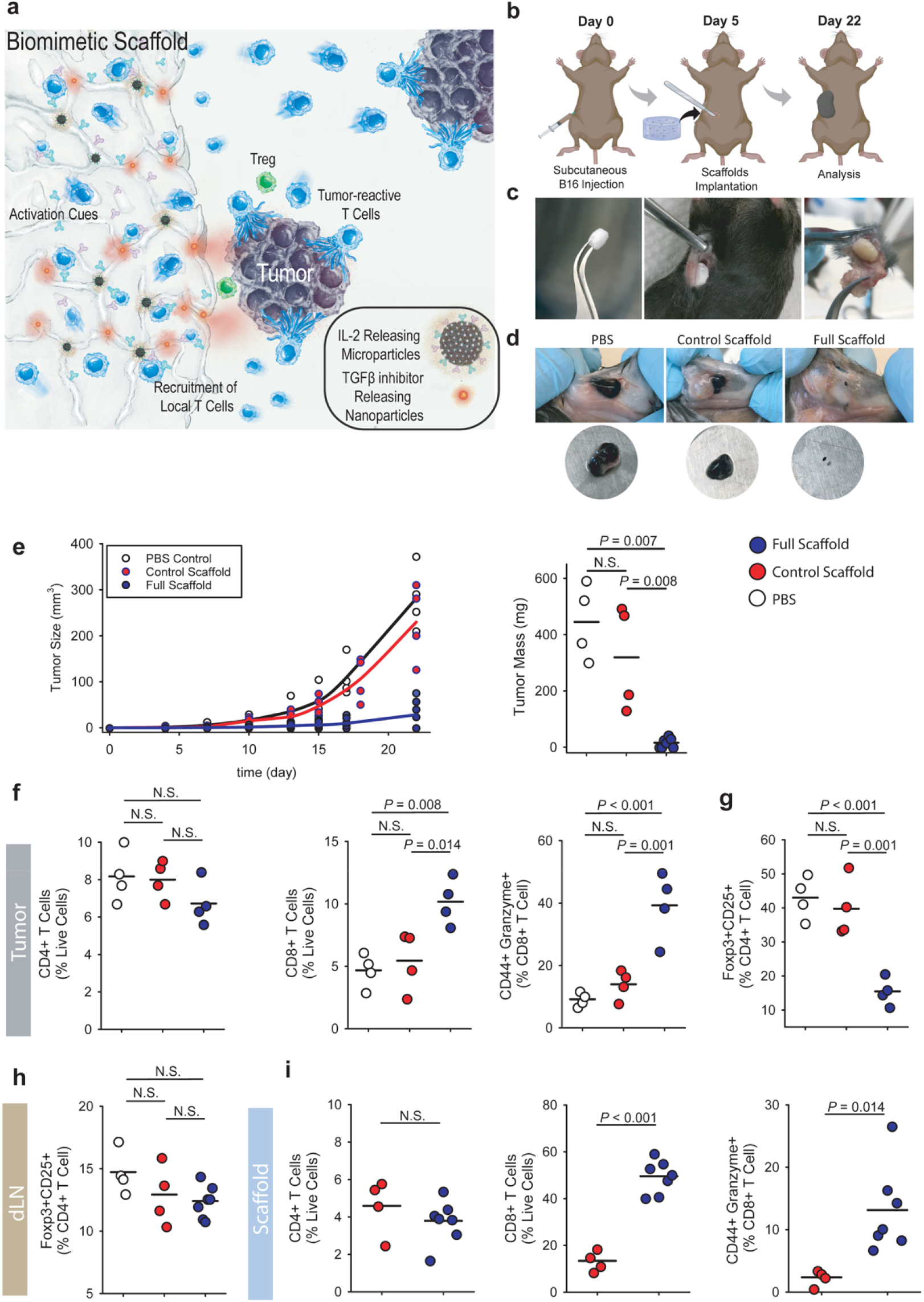
Engineered scaffolds that curb melanoma tumors. **(a)** Schematic representation of the scaffold components. Sustained release of CCL21 helps recruit endogenous T cells, while the presentation of surface-conjugated activation cues (anti-CD3 and anti-CD28) and the sustained release of IL-2 activates recruited T cells. Sustained release of TGF-β inhibitor will prevent formation of Tregs both in scaffolds and tumors. **(b)** Timing of tumor inoculation and follow up surgical implantation of the biomaterial scaffold. **(c)** The engineered device is surgically implanted in a B16-F10-ova bearing mice. **(d)** Melanoma (B16-F10-Ova) tumor growth and final tumor mass in wild-type mice implanted with either full (n = 7) or control scaffolds (n = 4) compared to PBS (n = 4). Each point represents one mouse. **(e)** Representative image of tumors extracted from wild-type mice 22 days after tumor inoculation. Local recruitments and activation of endogenous T cells plus Treg suppression via the implanted alginate-based scaffold successfully eliminated the aggressive melanoma tumor in mice. **(f)** *Status of recruited T cells in tumors*. Flow cytometry analysis of CD4+ and CD8+ T cells recruited and expanded in the tumors 22 days after subcutaneous injection of B16F10-ova cells. Activation of recruited CD8+ T cells was monitored by measuring surface expression of CD44 as well as intracellular measurement of Granzyme B expression. **(g**,**h)** The frequency of Foxp3+CD25+CD4+ Tregs in tumors and tumor-draining lymph nodes. **(i)** *Activated CD8+ T cells inside scaffolds*. Flow cytometry analysis of the percentage of CD8+ T cells in explanted scaffolds 17 days after subcutaneous implantation of scaffolds.

### B16-F10 melanoma model

To determine whether local depletion of Tregs could facilitate clearance of tumors, we tested our scaffolds in the murine melanoma model. Mice received a subcutaneous injection of B16-F10 cells to their right flank. We prepared scaffolds that were functionalized as above (“full”) and control scaffolds that lacked functionalization. After 5 days, when the tumors first became palpable, we implanted the scaffolds adjacent to the tumor in the subcutaneous space (**Fig. 1b-c**); some mice remained entirely untreated. Unfunctionalized scaffolds served as controls for the surgery and wound healing. We ended the experiment for all mice on day 22, when the untreated mice required euthanasia for tumor size and morbidity. H&E staining of the tissue adjacent to the tumor and scaffold showed successful tissue integration and recruitment of T cells *via* the implanted microporous scaffolds (**Extended Data Fig. 6a**). In all mice receiving the functionalized scaffold, tumor growth was suppressed (**Fig. 1d**) as compared with untreated mice or those receiving a control scaffold. The tumor size for treated mice was roughly one eighth the size of those receiving control scaffolds (**Fig. 1e**). For three (43%) of the treated mice, no residual tumor could be found at all (complete remission) (**Fig. 1e**).

Tumors, implanted scaffolds, tumors’ draining lymph nodes, and spleens were examined. The tumor-infiltrating T cells showed an increase in the proportion of CD8+ T cells, and they were highly activated and cytotoxic (**Fig. 1f**) in the tumors adjacent to functionalized scaffolds versus control scaffolds. These T cells were also significantly more activated and cytotoxic within the functionalized versus the control scaffolds (**Fig. 1i**). The proportions of total CD4+ T cells, on the other hand, were roughly consistent (**Fig. 1f**,**i**). The effect of recruitment and proliferation of the CD8+ T cells was local, as we saw no increase in the proportion, activation, or cytotoxicity of CD8+ T cells in the draining lymph node (**Extended Data Fig. 7a-c**). These findings show that the functionalized scaffolds successfully recruited and locally activated T cells, promoted their entry into tumors, and promoted the clearance of even aggressive tumors.

To better understand the impact of local TGF-β inhibition, we examined the tissues for the presence of Tregs. Tregs were depleted by about 60% in the tumors adjacent to functionalized scaffolds as compared with controls (**Fig. 1g**). As systemic administration of TGF-βi can result in autoimmune disease,^15^ we examined whether the local release of TGF-βi adjacent to the tumor could suppress Tregs in distant parts. We found no significant changes in Treg proportions in the draining lymph node (**Fig. 1h** and **Extended Data Fig. 7e**). Exhaustion of tumor infiltrating lymphocytes is commonly noted, and we saw no significant difference in PD-1^hi^ T cells in our tumors (**Extended Data Fig. 7d** and **8d**).

To better understand the mechanism of these underlying findings, we examined the T cells recruited to the scaffolds. Previous work in melanomas overexpressing CCL21 showed recruitment of both CD4+ and CD8+ T cells, albeit with different tempos^16^. The scaffolds above employed the chemokine CCL21 to recruit T cells into the milieu. We compared whether ligation of two different chemokine receptors on T cells, CCR7 and CXCR4, by two different chemokines, CCL21 and CXCL12 (also called SDF-1α), respectively, could explain the preferential recruitment of CD8+ T cells into the scaffold and tumor. Both chemokines successfully recruited T cells and helped clear tumors compared to control scaffolds (**Extended Data Fig. 9a-b**). We found that CCL21 favored recruitment of CD8+ T cells and disfavored CD4+ T cells as compared with CXCL12 (**Extended Data Fig. 9c**). Despite the presence of identical activation antibodies CD3 and CD28 in the scaffold, co-ligating CCR7 with CCL21 also increased the cytotoxic state of the effector T cells (**Extended Data Fig. 9d**). These results show that CCL21 improves recruitment of cytotoxic effectors into the tumor compared to CXCL12. Moreover, ligation of CCR7 improves the killing potential of CD8+ T cells. These results show that activation of chemokine receptors on T cells improves the therapeutic capability of our scaffold.

An important detail in the therapeutic translation of this scaffold lies is its stability prior to use. Water-based hydrogels can have low stability because of spontaneous hydrolysis, which we sought to reduce by preparing our alginate-based scaffolds in a sterile, lyophilized, and dry state. To test whether the scaffolds could retain their functional ability after 6 months of cold storage, we employed functionalized scaffolds in the B16 melanoma mouse model and found no difference between freshly fabricated scaffolds and older ones (**Extended Data Fig. 5** and **10**).

These results showed that the immunoactive scaffold cleared tumors by recruiting and highly activating CD8+ T cells while depleting local Tregs within the tumor.

### Distant tumors

By the time many solid tumors are clinically detected, they have already spread regionally or distantly. We next assessed whether local depletion of intratumoral Tregs in a primary tumor could alter the trajectory of the immune response in a distant secondary tumor. Wildtype mice received a subcutaneous injection of B16-F10 cells in the flank, and as before functionalized or unfunctionalized scaffolds were implanted subcutaneously after 5 days, when the tumors first became palpable. On the same day, we injected tumor cells contralaterally to primary one (**Fig. 2a**). We monitored tumor growth on both sides and measured tumor masses. The growth of the secondary tumor was suppressed by about 40 percent upon local treatment of the primary tumor with immunoactive scaffolds but not control scaffolds (**Fig. 2b-d**). The proportion of tumor-infiltrating CD8+ T cells was increased by more than two-fold in the contralateral tumor of the mice that received functionalized scaffolds (**Fig. 2e, Extended Data Fig. 11**). Activated T cells were also found in the spleen and draining lymph nodes, suggesting that local inhibition of Tregs allows widespread trafficking of activated effector T cells (**Extended Data Fig. 13**,**14**). Those T cells were more activated and expressed more Granzyme B (**Fig. 2f** and **Extended Data Fig. 12c-d**). We noted the expansion at 22 days of an expanded population of CD62L+CD44+ CD8+ T cells, suggesting the development of central memory T cells in the tumor. This T cell memory subset has been noted in mice and humans to be important for both clearance of tumors and prevention of their recurrence^17^. As before, the proportion of Tregs was markedly reduced in the primary tumor. Importantly, Tregs were also reduced in the secondary tumor, but to a lesser extent (**Fig. 2h,i**), but not in the lymph nodes (**Extended Data Fig. 13e**). These results show that localized suppression of intratumoral Tregs resulted in both successful clearance of primary tumors as well as marked reduction of secondary tumors. The minor decrease in Tregs in the distant tumor supports the model that antigen-specific Tregs trained in the primary tumor may spread globally to facilitate distant tumors. By suppressing those Tregs in the primary tumor, we suppressed Tregs in the secondary tumors. However, the disproportionate increase in activated, cytotoxic CD8+ T cells in the secondary tumor despite the minor decrease in Tregs there indicates that expansion of potent effector T cells in the primary tumor spreads to secondary sites, an immunological “abscopal” effect on distant sites arising from the scaffold acting on the primary tumor.

**Fig. 2.**
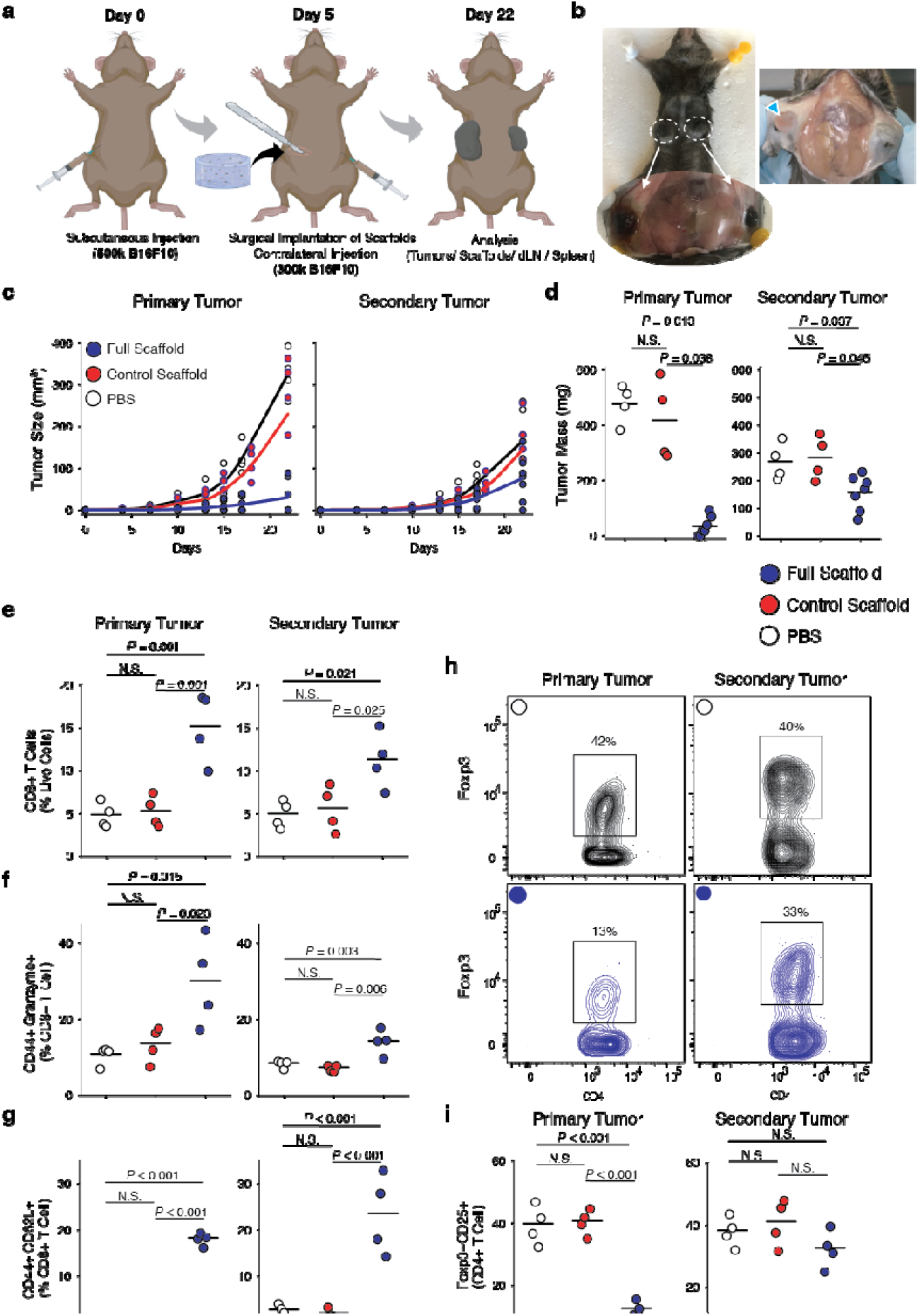
Engineered scaffolds can suppress the growth of not only local tumors but also distant tumors. **(a)** Timing of inoculation of primary and secondary tumors and follow up surgical implantation of the cell-free scaffolds. **(b)** Growth of primary and secondary tumors in the control mouse. **(c)** Melanoma (B16-F10-Ova) tumor growth and **(d)** final tumor masses were measured 22 days after inoculation of primary tumors in wild-type mice treated with Full or control scaffolds (n= 4-7). Each point represents a mouse. **(e)** The frequency of CD8+ T cells in primary and secondary tumors (n=4). Flow cytometry study of the frequency of **(f)** CD44+GZMB+CD8+ (activated) and **(g)** CD44+CD62L+CD8+ (central memory) T cells in primary and secondary tumors (n=4). **(h)** Representative flow cytometry of Foxp3+CD25+CD4+ Tregs in primary and secondary tumors for mice treated with Full Scaffolds (Blue) and PBS (Black). **(i)** The quantified frequency of Foxp3+CD25+CD4+ Tregs in primary and secondary tumors.

### Adoptive T cell therapy

Delivery of engineered T cells like CAR-T cells is necessary for a variety of cancer therapies, but unlike in lymphomas, intravenous administration does not always allow for efficient delivery of tumor-specific T cells into solid tumors^18^. Alginate-based scaffolds that lack structure have been used to deliver T cells in murine melanoma and breast cancer^14,19^, where the matrix served as a T-cell depot to support their local expansion. We sought to employ the structural advantages of microporosity and mechanical rigidity of our scaffold to deliver T cells. Wildtype mice received a subcutaneous injection of B16-F10 cells in the flank, and as before functionalized or unfunctionalized scaffolds were implanted after 5 days, when the tumors first became palpable (**Fig. 3a-d)**. These scaffolds were loaded just before implantation with activated OT-I T cells. Some mice instead received intravenous injection of either the same number of OT-I T cells or saline (**Fig. 3d-f** and **Extended Data Fig. 17**). Mice were euthanized at 22 days as before for further analysis.

**Fig. 3.**
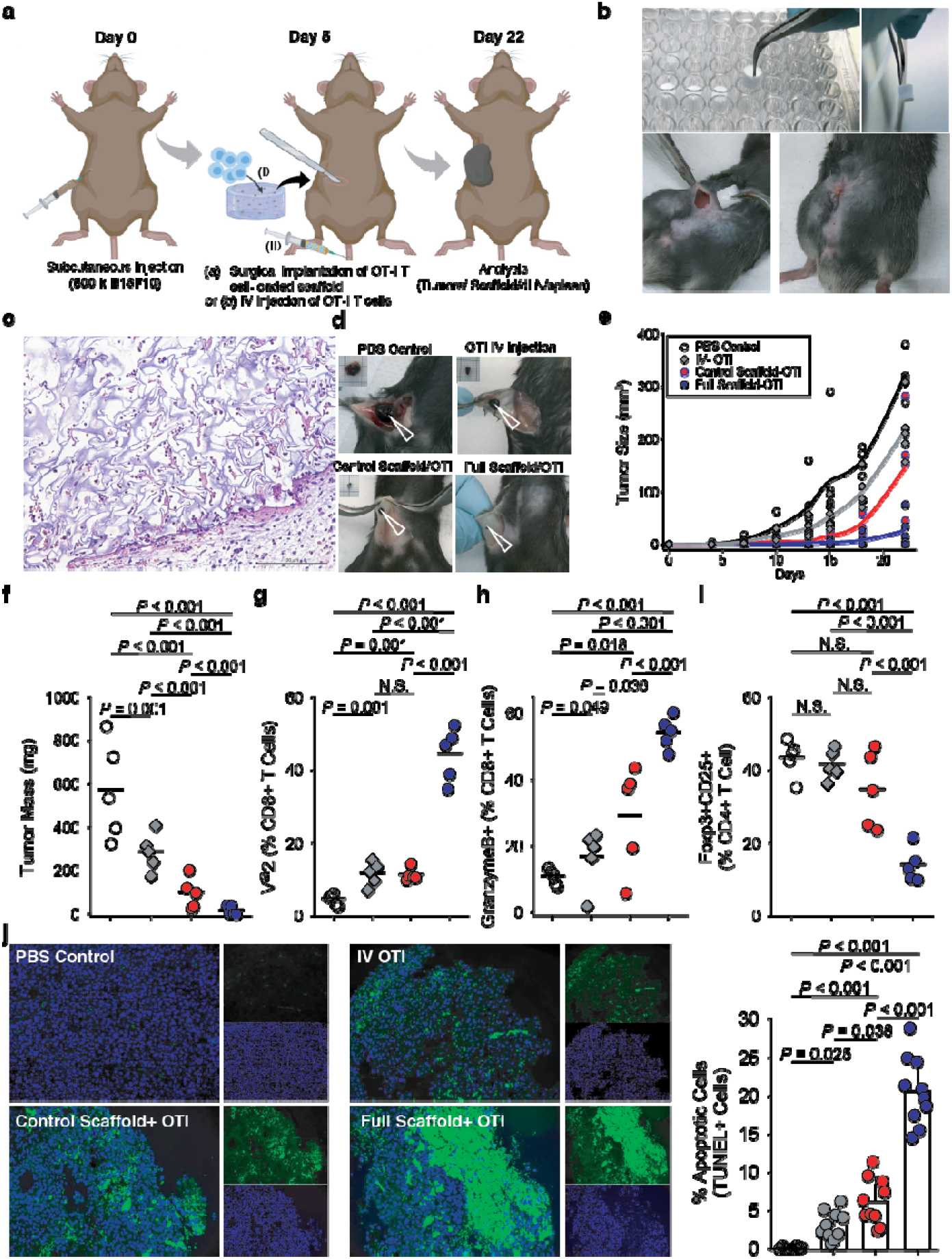
Engineered scaffolds deliver antigen-specific T cells and suppress the growth of melanomas. **(a)** Timing of tumor inoculation and implantation of the activated OTI cells-loaded scaffold. **(b)** The engineered device is surgically implanted in a B16-F10-ova bearing mouse. **(c)** H&E staining showing connective tissue-like scaffolds. **(d)** Representative images of subdermal tumors from wild-type mice 22 days after tumor inoculation. Alginate scaffolds carrying tumor reactive T cells and T cell-specific activator cues can eliminate melanoma tumors in mice. **(e)** Melanoma (B16-F10-Ova) tumor growth and **(f)** final tumor mass in wild-type mice with immunoactive or control scaffolds compared to PBS (n=5). Each point represents one mouse. **(g)** Presence of tumor specific CD8+ T cells (OTI) in tumors was studied 22 days after inoculation of tumor cells using flow cytometry. **(h)** Frequency of CD8+ T cells with high expression of GZMB effector cytokine. **(i)** The frequency of Foxp3+CD25+CD4+ Tregs in tumor were studied (n= 5). **(j)** Tumor-cell apoptosis was detected using TUNEL staining after various treatments, examples of staining on the left and summarized data on the right.

H&E staining of the scaffolds adjacent to the tumor confirmed tissue engagement, successful delivery, and proliferation of OT-Is plus recruitment of endogenous T cells (**Fig. 3c**). Our OT-I-loaded, functionalized scaffold significantly suppressed tumor growth compared to other treatments and controls, by ∼16-fold compared to saline control and ∼10-fold compared to IV injection of OT-I T cells (**Fig. 3e-f** and **Extended Data Fig. 15**). These results show that scaffold-based delivery resulted in better tumor clearance than intravenous delivery of T cells.

Moreover, we found that the accumulation and activation of antigen-specific T cells was far more effective with the immunoactive scaffolds than with delivery by the inert, control scaffold or intravenous delivery (**Fig. 3g-i**). Scaffold-based delivery of OT-I T cells dramatically outperformed intravenous delivery when looking at the resulting number of OT-I T cells in the tumor (**Fig. 3g**). Moreover, the scaffold-delivered T cells were more activated and showed greater cytotoxicity (**Fig. 3h** and **Extended Data Fig. 16**). Treg populations in the tumor showed significant differences between treatment groups. Regulatory T cells were suppressed by over 60 percent in the tumors adjacent to immunoactive scaffolds as compared to the control scaffold, intravenous delivery of T cells, or saline controls (**Fig. 3i**). TUNEL staining showed a significant increase in apoptotic tumor cells in the functionalized scaffold that delivered OT-I T cells versus controls (**Fig. 3j**). These results confirm that the immunoactive scaffold delivers highly cytotoxic, antigen-specific T cells to the tumor environment.

Tumor-draining lymph nodes showed a comparable number of OT-I T cells when they were delivered by the scaffold as compared with IV injection (**Extended Data Fig. 17**). On the other hand, highly activated OT-I cells were more present in the LN and spleens when mice received an IV injection versus when scaffolds delivered the T cells (**Extended Data Fig. 17** and **18)**. As before, the proportion of Tregs in lymph nodes were comparable between groups (**Extended Data Fig. 17c**). These results explain why intravenous injection of engineered, tumor-specific T cells so often fails: these activated T cells end up in the LNs and spleen rather than in the tumor. The direct access of effector T cells to the tumor microenvironment provided by the scaffold also bypasses the formidable barriers established by the tumor vasculature to the transport of tumor-specific T cells^20^.

In summary, we demonstrate that an approach orchestrating multiple immunological mechanisms simultaneously can overcome barriers imposed by solid tumors to allow endogenous or engineered T cells to infiltrate and eradicate otherwise aggressive cancers.

## Online Methods

### Experimental Materials and Methods

#### Chemicals and Biologicals

Unless noted otherwise, all chemicals were purchased from Sigma-Aldrich, Inc. (St. Louis, MO). All glassware was cleaned overnight using concentrated sulfuric acid and then thoroughly rinsed with Milli-Q water. All the other cell culture reagents, solutions, and dishes were obtained from Thermo Fisher Scientific (Waltham, MA), except as indicated otherwise.

#### Preparation and characterization of cytokine-producing microparticles

Monodisperse mesoporous silica microparticles (5 to 20 μm) were formed using a microfluidic jet spray-drying process, using cetyltrimethylammonium bromide (CTAB) and/or Pluronic F127 as templating agents, and tetraethylorthosilicate (TEOS) for silica as reported before.^21,22^ To conjugate heparin to the particles, we first added amine groups to the particles. We suspended the mesoporous silica microparticles (800 mg) in dehydrated methanol (50 mL) then added APTES (3 mL) and stirred at room temperature overnight. The final product was centrifuged (1500 rpm, 3 min) and washed with methanol five times, followed by drying under high vacuum. To functionalize the aminated-silica particles with heparin, heparin sodium salt (216 mg) was dissolved in deionized water (8 mL) and activated via successive addition of EDC (63 mg) and sulfo-NHS (71.4 mg). After stirring for 5 min, the ethanolic solution of amino-functionalized silica (20 mg in 1.12 mL) was added to the reaction mixture and stirred for 12 h at room temperature. Afterwards the particles were separated by centrifugation and washed several times with deionized water and ethanol to remove unreacted reagents.

To incorporate IL-2 (Interlukine-2, BRB Preclinical Repository, NCI, NIH, Frederick, MD) into these heparin-coated microparticles, the microparticles were incubated with different amounts of cytokine in PBS buffer containing bovine serum albumin (BSA; 0.1 %w/v) and were gently shaken overnight at 4°C. The microparticles were then centrifuged and washed several times to remove unabsorbed cytokines.

To estimate the *in vitro* release of IL-2 from microparticles, we incubated 20 × 10^6^ microparticles in 2 mL PBS (pH 7.4; supplemented with 1 mM CaCl_2_) at 37 °C. At different time intervals, 500 µL of the supernatant was collected, and replaced with an equivalent volume of PBS. The concentration of released IL-2 was determined using a human IL-2 ELISA kit (ThermoFisher human IL-2 Kit). The final particles used for formulating scaffolds (below) employed 50 μg of IL-2 per mg of particles.

The silica particles were also covalently conjugated with T-cell activating antibodies: anti-CD3 (clone 2C11; Bio-X-Cell) and anti-CD28 (clone 37.51; Bio-X-Cell). We used 1 μg anti-CD3 and 0.25 μg anti-CD28 per 10^6^ silica particles. After activation of antibodies’ carboxylic groups for 10 min with EDC/NHS, microparticles were added and incubated under gentle stirring at 4 °C overnight. The functionalized microparticles were then separated from the solution and washed several times. Unreacted functional groups were quenched by washing samples in Tris buffer (100 mM, pH 8) for 30 min. Micro-BCA assay was used to quantify total amount of surface conjugated antibodies according to the manufacturer’s protocol.

#### TGF-β inhibitor particles

We prepared poly(lactic-co-glycolic) acid (PLGA) nanoparticles (NPs) to release the TGF-β inhibitor LY2157299 (also called galunisertib, Cayman Chemical). We prepared the PLGA using a nanoprecipitation method as previously reported^23^: briefly, Resomer RG 503 PLGA (50:50; molecular weight: 28 kg/mol) was used in this study. LY2157299 and PLGA were dissolved in 5 mL dichloromethane and sonicated into 1% poly vinyl alcohol (PVA) solution (50 mL) by probe sonicator (12 W) for 2 min. The resulting emulsification was then added to 100 mL of 0.5% PVA solution. The solution was agitated, and the dichloromethane was allowed to evaporate for 4 h. The solution was then centrifuged at 3000 × *g* for 5 min to pellet out any non-nano size material. The supernatant was removed and ultracentrifuged and washed three times at 21,000 g for 20 min to wash away the PVA. The resulting nanoparticle solution was flash frozen in liquid nitrogen and lyophilized for 2 days prior to characterization and use. Hydrodynamic diameter and surface charge of formed PLGA NPs were studied using dynamic light scattering (DLS) and zeta potential measurements (Zetasizer Nano, Malvern, UK).

#### Preparation and characterization of scaffolds

To form the scaffolds, we first oxidized the alginate (Mw ∼250 kDa, high G blocks; Novamatrix UP MVG, FMC Biopolymer, Rockland, Maine) with sodium periodate (1.5 %), overnight at room temperature, then quenched the reaction by dropwise addition of ethylene glycol for 45 min. We then dialyzed the solution (MWCO 3.5 kDa) against deionized water for 3 days followed by lyophilization. Afterward, the alginate was dissolved in MES (MES 150 mM, NaCl 250 mM, pH 6.5) and covalently conjugated to RGD-containing peptide (GGGGRGDY; GenScript USA Inc., Piscataway, NJ) using carbodiimide chemistry (NHS/EDC). The reaction was continued for 24 h followed by dialysis (MWCO 20 kDa) and lyophilization. This alginate-RGD complex in PBS was mixed with CCL21 protein (Peprotech Recombinant Murine Exodus-2, 250 ng per scaffold). To prepare scaffolds, 20×10^6^ IL-2-loaded heparin-functionalized silica microparticles were mixed with 1 mL of alginate. The alginate / chemokine / cytokine-particle / TGF-βi-NP mixture was cross-linked using calcium sulfate solution. Each resulting scaffold bore 400 IU of IL-2. The gels were casted in desired 24- or 96-well plates followed by two overnight washes to remove extra calcium ions. These scaffolds were frozen at -80°C, lyophilized for 3 days, and stored at 4 °C before use.

To load these NPs into alginate-based scaffolds, LY2157299-loaded PLGA NPs were mixed with alginate prior to crosslinking *via* calcium. The concentration of released LY2157299 from nanoparticles before and after loading into alginate scaffolds was determined by measuring the UV absorption of LY2157299.

We prepared an array of different alginate formulations by varying either the polymer content or the amount of crosslinker (here CaSO_4_). To measure the mechanical stiffness of our gels we used an Instron 5542 mechanical tester and all the samples were tested at a rate of 1 mm/min. The Young’s modulus was then calculated from the slope of the linear region that corresponds with 0–10% strain. The final alginate formulation employed for in vivo studies comprised alginate 2.5% with 40 mM CaSO_4_.

To sterilize the fabricated scaffolds before *in vitro* or *in vivo* functional assays, we employed X-ray irradiation (Gulmay Medical RS320 X-ray unit) as per ISO 11137-2:2013 recommended protocols.^24^ A dose of 25 kGy (2.5 Mrads) was used for sterilization. Physical properties, including changes in morphology and mechanical stiffness of the scaffolds, or T-cell activation property change after sterilization was tested (**Extended Data Figure 6D**,**E**).

Scanning electron microscopy (SEM) images of the gels were taken to see the cross-sectional microstructure and porosity of the alginate-based scaffolds. The lyophilized scaffolds were freeze-fractured (using liquid nitrogen) for cross-sectional images. The scaffolds were sputtered with iridium (South Bay Technology Ion Beam Sputtering) prior to imaging with a ZEISS Supra 40VP scanning electron microscope (Carl Zeiss Microscopy GmbH). The sizes of pores from different parts of the SEM images were then measured and analyzed using ImageJ software (NIH). For SEM imaging of cell-loaded scaffolds, the cell-laden hydrogels were fixed with 2.5% glutaraldehyde, followed by post-fixation in osmium tetroxide prior to serial dehydration in increasing concentrations of ethanol (25, 50, 75, 90, and 100%) for 15 min each, and iridium sputtering.

To immobilize anti-CD3 and anti-CD28 to the scaffolds, the freeze-dried scaffolds were activated with EDC/NHS for 15 min. Then the scaffolds were washed twice with PBS (supplemented with 0.42 mM CaCl_2_) before addition of anti-CD3 and anti-CD28. Then they were incubated at 4°C overnight. Unreacted functional groups were quenched by washing the scaffolds with Tris buffer (100 mM, pH 8) for 30 min. For T cell activation studies, 5 × 10^6^ primary naïve T cells were added to the scaffolds and cultured for 3-5 days to study their effector functions.

CCL21 release from the scaffolds was measured by murine CCL21 ELISA kits as a function of time.

#### T-cell isolation and activation

All *in vitro* experiments were conducted in accordance with UCLA’s institutional policy on humane and ethical treatment of animals following protocols approved by the Animal Research Committee. Five- to eight-week-old wild-type or OT-I TCR transgenic mice (Jackson Labs) were used for all experiments.

T cells were purified from spleens using the EasySep immunomagnetic negative selection enrichment kit (Stem Cell Technologies). T cells were cultured in media comprising RPMI-1640 supplemented with 10% heat-inactivated FBS, 1% penicillin/streptomycin, 1% sodium pyruvate, 1% HEPES buffer, 0.1% μM 2-mercaptoethanol.

Control *in vitro* activation of T cells was performed by culturing 1 × 10^6^ cells/mL in tissue culture-treated 24-well plates that were pre-coated with anti-CD3 (clone 2C11; Bio X Cell) at a concentration of 10 μg/mL plus addition of 2 μg/mL soluble anti-CD28 (clone 37.51; Bio X Cell). T cells were then collected from wells and allowed to proliferate in interleukin-2 (IL-2, BRB Preclinical Repository, NCI, NIH)-containing medium (50 U/mL), prior to being used for experiments.

For Treg formation experiments CD4+ T cells were purified from mouse spleen as mentioned above. Cells were then either activated on scaffolds or on anti-CD3e antibody (8 mg/ml) coated plates with the anti-CD28 antibody (2 mg/ml) supplemented medium. At the same time TGF-β (15 ng/ml) was added to the media. After four days regulatory T cells were removed and stained with antibodies for flow cytometry analysis.

#### Flow cytometry

For flow cytometry analysis, antibodies to mouse antibodies, were purchased from eBioscience, BioLegend, or BD Biosciences. To study proliferation behavior of T-cell responses during various treatments their expansion was measured by 5-(and-6)-carboxyfluorescein diacetate, succinimidyl ester (CFSE) dilution. For CFSE dilution experiments, 5 × 10^5^ naive CD4+/CD8+ T cells were labeled with 2 μM CFSE for 13 min, followed by two washes and then incubation with splenocytes. Splenocytes were extracted from the spleens of wild type C57Bl/6 mice. Then the cells were incubated in ACK lysis buffer (Gibco) for 5 min at room temperature to remove red blood cells. The remaining cells were then treated with ova peptide as above to present to T cells. Cells were analyzed on a Cytek DxP10 flow cytometer using FlowJo software (Treestar/BD).

For intracellular staining of GranzymeB and Foxp3, we followed the manufacturer’s protocol (Foxp3 / Transcription Factor Staining Buffer Set, eBioscience). The following antibodies were used for intracellular staining from Biolegend: Foxp3 (clone MF-14, AF647, Cat #126408); GranzymeB (clone GB11, AF647, Cat #515406), Mouse IgG1, κ Isotype Ctrl (clone MOPC-21, AF647, Cat #400130).

#### Migration Assay

The migration assay to evaluate the role of chemokines on recruitment of T cells and melanoma cells in the presence and absence of chemokine-functionalized scaffolds was performed using regular Transwell migration as we reported.^9^ The number of migrated cells was evaluated after 4 h using an automatic cell counter.

#### Tumor Assays

2-5 × 10^5^ B16F10-OVA tumor cells were subcutaneously injected into right or both (in the contralateral tumor model) right and left flanks of C57BL/6J WT mice (6-8 weeks old). These melanoma-derived cells are transfected to express chicken ovalbumin peptide (OVA)^25^. Five days after tumor cell injection, scaffolds were surgically implanted subcutaneously into the same approximate region of the tumors in both flanks. For cell-loaded studies, *ex vivo* activated OT-I T cells were transferred either intravenously using retro-orbital injections (100 µL per animal) or implantable scaffolds at the same day. Tumor size was assessed over time using a digital caliber until day 22 at which animals were sacrificed and the tumor, draining lymph nodes, and spleen were extracted. Tumor mass was measured using a digital balance before digesting the tumor tissue for flow cytometry or fixing it for tissue sectioning. Tumors were digested by incubating in collagenase and DNase I (50 μg/mL) at 37 °C for 15 min. These enzymes were inactivated with EDTA (20 μL/mL of solution). Tissues were then mechanically disaggregated and passed through a 0.7 μm cell strainer to obtain a single-cell suspension. Cells were then stained with the fluorochrome-conjugated antibodies on ice. For intracellular staining, cells were permeabilized with Granzyme B Fix/Perm buffer according to the manufacturer’s instructions (BioLegend) before staining. Detection of apoptotic cells in tumor tissue was achieved using Terminal deoxynucleotidyl transferase-mediated dUTP nick-end labeling (TUNEL) staining following the manufacturer’s directions. TUNEL-positive cells indicated as apoptotic melanoma cells. Tissue sections were imaged by a fluorescence microscope (Keyence BZ-X800, Osaka, Japan).

#### Statistical analysis

The Kruskal-Wallis rank sum test, one-way ANOVA and two-tailed Student’s t-test were utilized as appropriate to analyze the data at a significance of α or p < 0.05. Quantitative data were expressed as mean ± standard deviation (SD). To determine the number of specimens for the proposed experiments, power analysis was conducted based on our preliminary data.

## Notes

### Competing Interest Statement

The authors have declared no competing interest.

