## Supplemental Figures for "Global anti-tumor immunity after localized, bioengineered Treg depletion"

### Extended Data

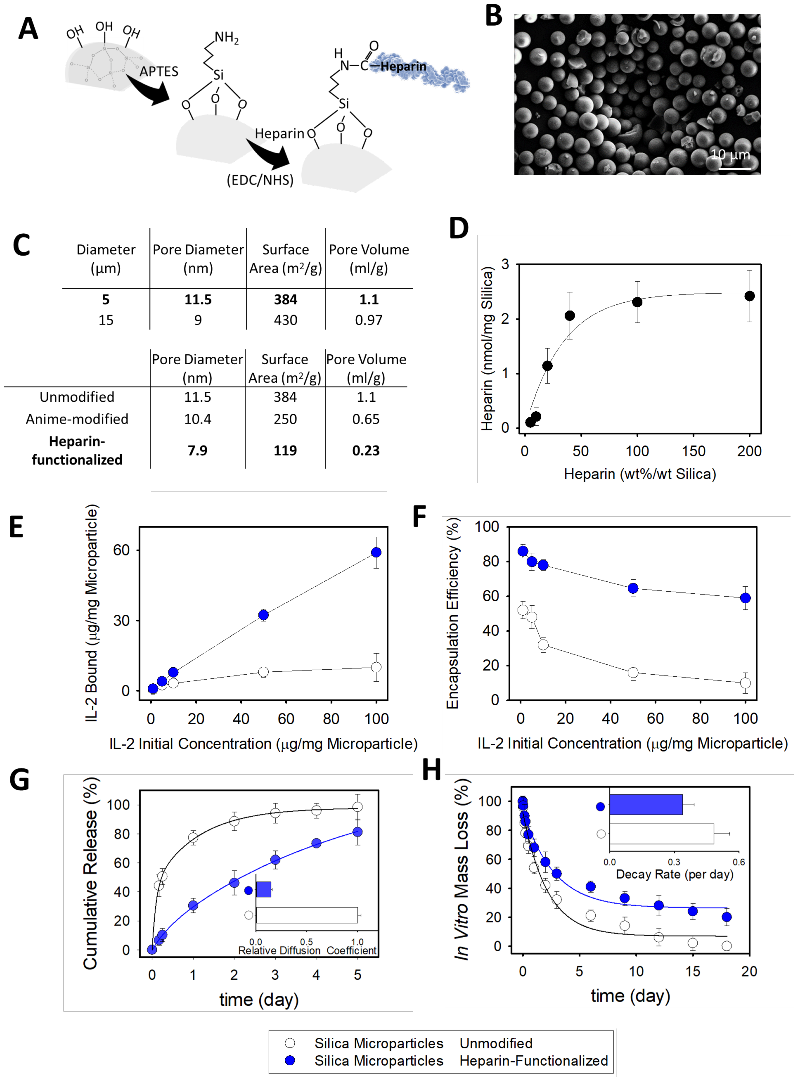


**Extended Data Fig. 1 | IL-2 cytokine is released from heparin-coated mesoporous silica microparticles**. **(a)** Surface chemistry for conjugation of heparin. **(b)** SEM image of synthesized mesoporous microparticles. **(c)** Change in physical characteristics of mesoporous silica microparticles after surface functionalization with APTES and heparin. **(d)** The degree of heparin-conjugation of silica particles with various initial amounts of heparin in the reaction mixture. **(e)** Binding efficiency of IL-2 to the unmodified compared to heparin-functionalized microparticles. **(f)** Encapsulation efficiency of IL-2 in unmodified compared to heparin-functionalized microparticles as a function of initial IL-2 concentration. **(g)** Cumulative release of IL-2 from heparin-functionalized and unmodified silica microparticles at 37 °C. (inset) calculated diffusion coefficients. **(h)** In vitro degradation of silica-based microparticles over time. (**e-h**) unmodified (open circles) compared to heparin modified (blue-filled circles) microparticles.

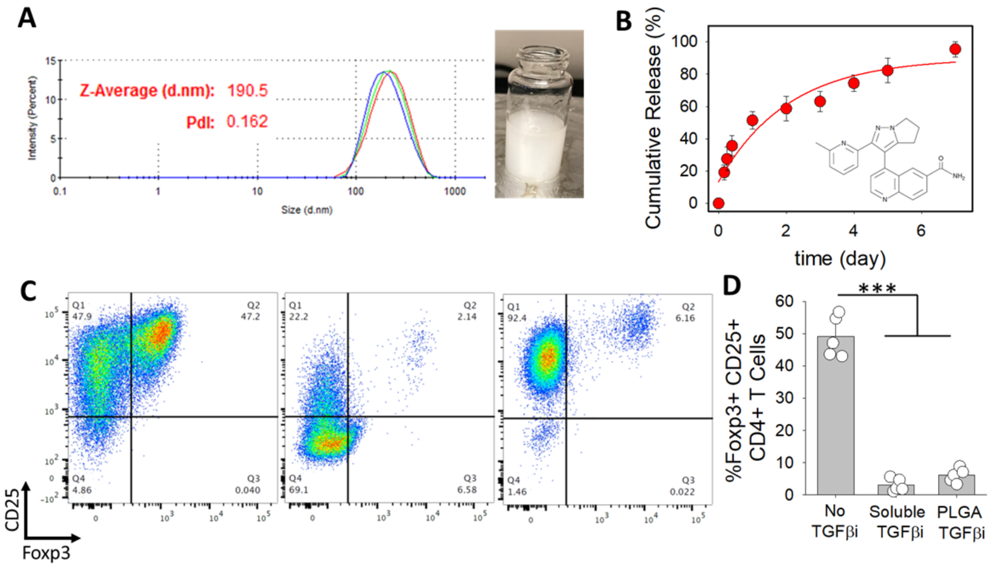


**Extended Data Fig. 2 | PLGA nanoparticles loaded with TGF-βi.** **(A)** Dynamic light scattering (DLS) showing monodisperse formation of TGF-βi-loaded PLGA nanoparticles. Inset showing stable suspension of formed nanoparticles in water 24 h after dispersion. **(B)** Release of TGF-βi from nanoparticles over time at 37 °C. Chemical structure of selected TGFbi, LY2157299, was also shown. **(C)** Left- Activation of naïve helper T cells results in formation of Tregs, using anti-CD3/28-coated aAPCs in the presence of soluble TGF-β. Middle- Inhibition of Treg formation using soluble TGF-βi (10 μM) or PLGA NPs loaded with equivalent amounts of TGF-βi. **(D)** Quantified percentages of formed Tregs.

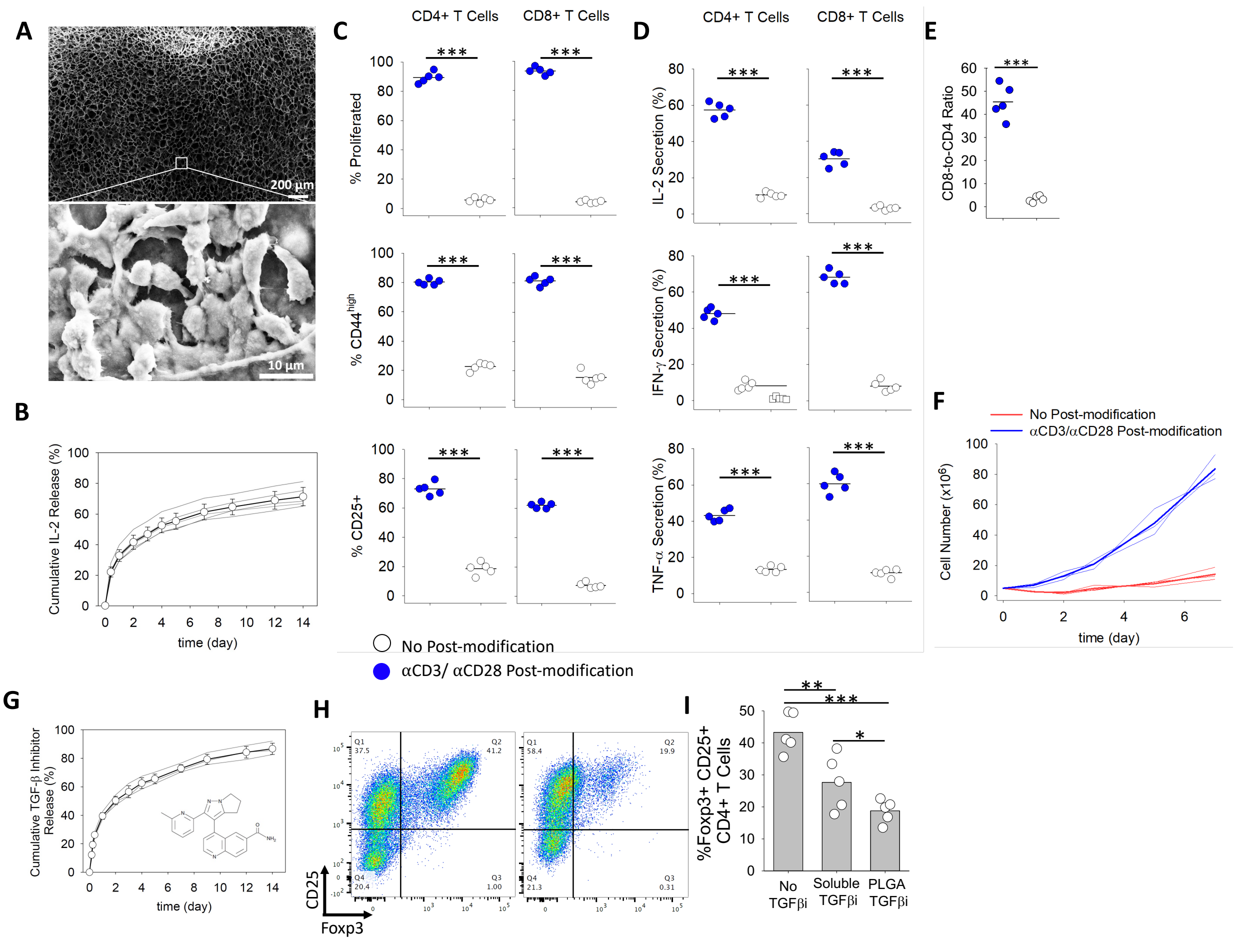


**Extended Data Fig. 3 | In vitro T-cell activation in 3D scaffolds.** (**A**) SEM images of macroporous 3D scaffolds. Images were taken from a region within the bulk of the scaffold. Scale bar is 200 µm. Enlarged SEM images demonstrating association of T cells within the alginate-based scaffolds. Images were taken from a pore wall of the scaffold where T cells were aligned. Scale bar is 10 µm. **(B)** Release of IL-2 encapsulated within mesoporous silica microparticles embedded in the alginate 3D scaffold. The released IL-2 was measured by ELISA over time under gentle shaking (50 rpm) at 37 °C. **(C-D)** *T-cell activation is modulated by aAPCs-loaded 3D scaffolds.* **(C) Activation and proliferation.** Naïve CD4+ and CD8+ primary mouse T cells co-cultured with 3D scaffolds. Scaffolds formulated by surface coating with stimulatory antibodies (anti-CD3 and anti-CD28) resulted in robust responses. Flow cytometry analysis of cell division (CFSE dilution) and expression of activation markers CD25 or CD44 assayed three days after the introduction of T cells into the biomaterial. %Proliferated is the percentage of T cells that divided at least once. Percentage of T cells with high expression of CD44 and percentage of T cells upregulating CD25. **(D) Cytokine production**. Naïve CD4+ and CD8+ primary mouse T cells co-cultured with 3D scaffolds. Scaffolds formulated by surface coating with stimulatory antibodies (anti-CD3 and anti-CD28) resulted in robust responses. Percentage of T cells expressing the effector cytokines IL-2, IFN-γ, or TNF-α. Each dot represents one independent experiment. **(E)** FACS quantification of CD8-to-CD4 ratio of T cells cultured with varying formulations of particles, compared to Dynabeads. The starting ratio for all conditions was 0.5 (two CD4+ T cells for each CD8+ T cell). **(F)** Naïve CD4+ and CD8+ T cells co-cultured with various formulations of 3D scaffolds. The starting number of T cells was $5x10^6$ cells. Absolute counts of viable T cells in scaffolds fabricated with different formulations shows the robust expansion of T cells. Each line represents an independent experiment. **(G)** Release of TGF-βi encapsulated PLGA nanoparticles embedded in the 3D alginate scaffold. The released TGF-βi was measured over time under gentle shaking (50 rpm) at 37 °C. **(H)** Flow cytometry analysis of Treg formation using antigen-presenting scaffolds in the absence (left) and presence (right) of TGF-βi releasing nanoparticles 4 days into culture. **(I)** Inhibition of Treg development in 3D scaffold by soluble TGF-βi versus PLGA-encapsulated TGF-βi was quantified. The individual data are presented (n=5). The results were statistically analyzed using one-way ANOVA with post-hoc analysis.

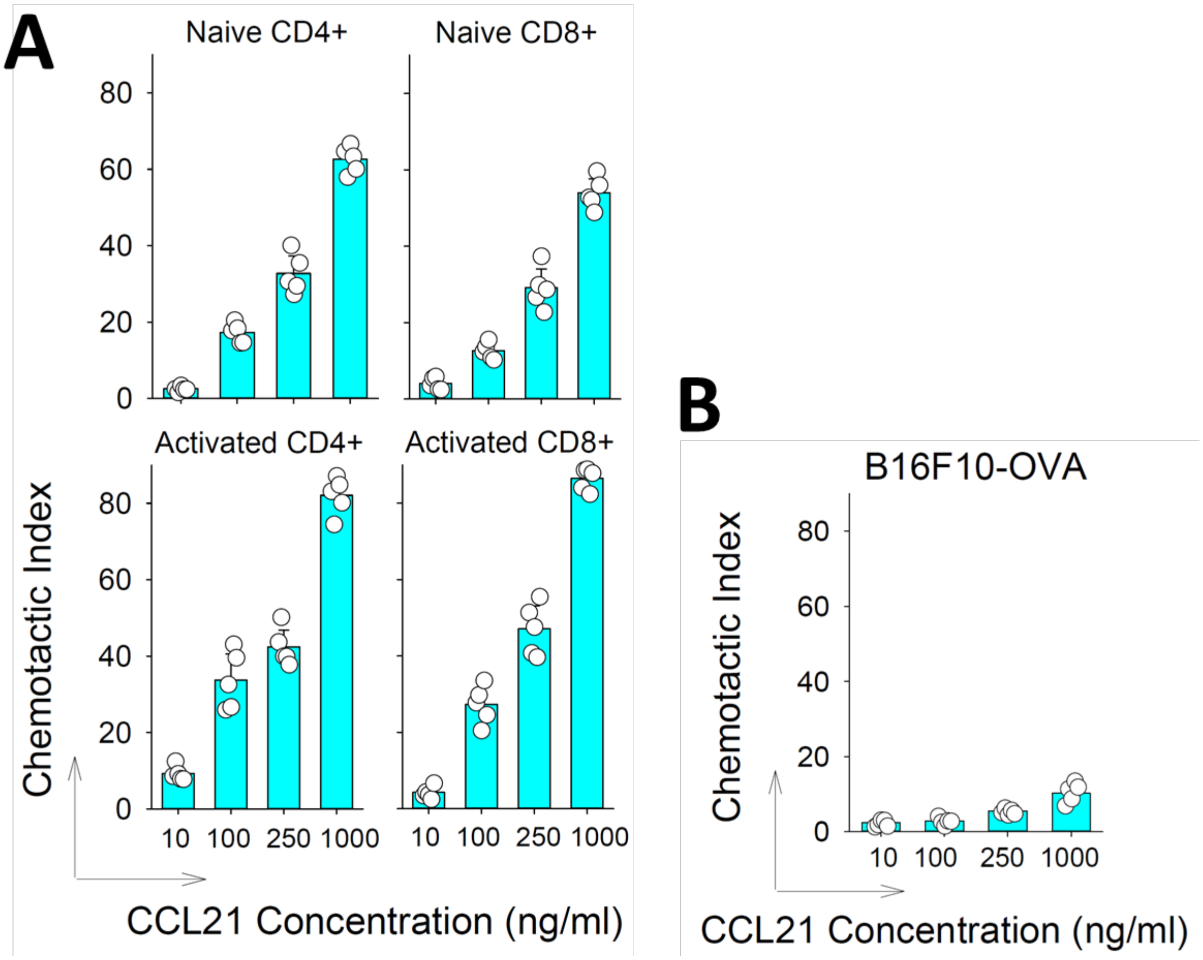


**Extended Data Fig. 4 | Assessment of CCL21 chemotaxis in recruitment** of **(A)** naive and activated CD4+ and CD8+ T cells and **(B)** B16F10-OVA tumor cells *in vitro*. $5\times10^5$ naive or activated T cells were loaded on the top filter of the transwell chamber. Hydrogels containing various concentrations of recombinant CCL21 were placed in the bottom wells at the indicated concentrations. Viable cells migrating to the lower chamber after 4 h (T cells) or 8 h (B16 cells) were quantified after digesting the scaffold. Chemotactic Index: fold migration over background (empty scaffolds). 5 μm pore size was selected for Transwell migration assay.

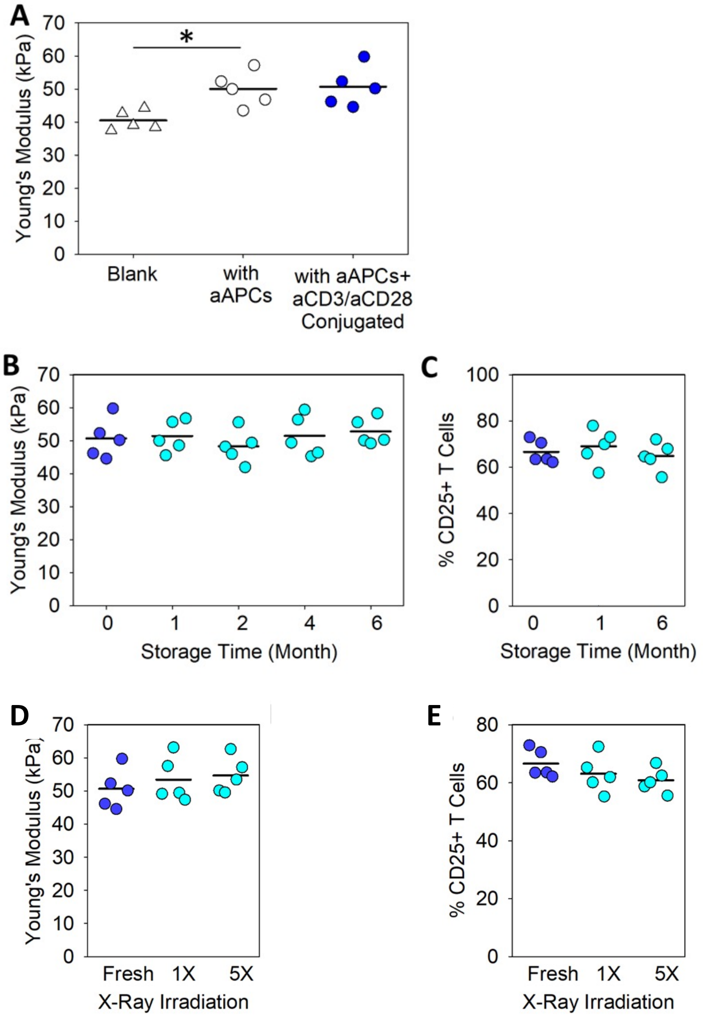


**Extended Data Fig. 5 | Mechanical characteristics of the 3D scaffolds. (A)** Change in mechanical properties of 3D alginate scaffolds in absence or presence of IL-2 releasing silica-based aAPCs with or without surface coating of the scaffold with anti-CD3/ anti-CD28 antibodies. **(B-C) Shelf-life evaluation of scaffolds.** Freeze-dried hydrogel batches were stored at 4 °C for different durations up to six months and changes in (**B**) mechanical properties were assessed as in A. (**C**) Assessment of activation capability of the 3D scaffold after storage. Scaffolds were prepared with microparticles comprising IL-2-releasing silica microparticles with and without surface coating of the scaffold with anti-CD3/ anti-CD28 antibodies and stored for various durations. Scaffolds were then used for co-culture with naïve, primary CD8+ T cells. Activated T Cells were assessed by upregulated expression of CD25. **(D)** Change in mechanical properties and **(E)** T cell activation of scaffolds after 1 or 5 cycles of x-ray irradiation at 25 kGy dose compared to freshly prepared samples. Lyophilized scaffolds were exposed to X-ray exposures with 24 h time intervals. **(A-E)** Each dot represents an independent experiment (n=5). The results were statistically analyzed using one-way ANOVA with post-hoc analysis. In (B-E) Results showed no statistically significant ($p > 0.05$) changes in elastic modulus or T-cell activation.

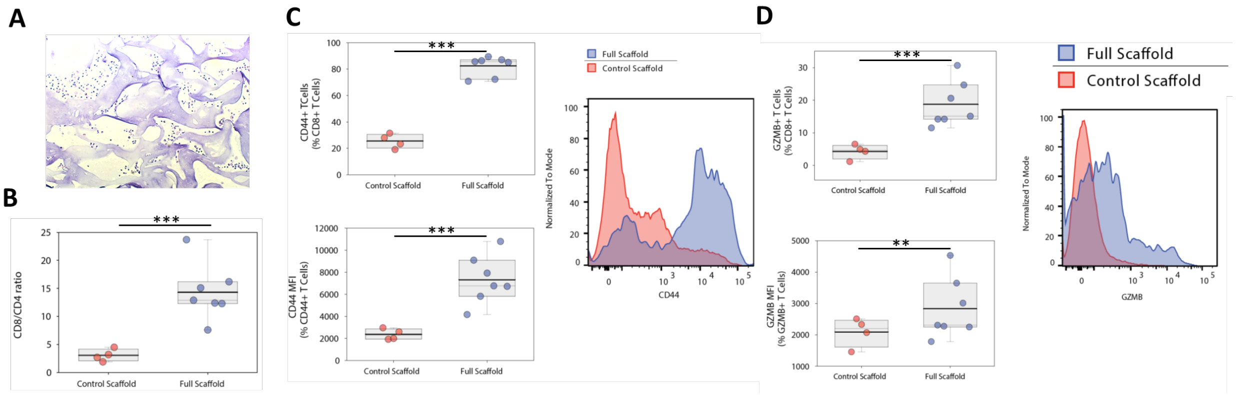


**Extended Data Fig. 6 | Activation of endogenous T cells recruited <u>into scaffolds</u>. (A)** H&E staining of the cross sections of the subcutaneously implanted scaffolds that originated from the alginate biopolymer, 7 days after implantation. **(B)** FACS quantification of CD8-to-CD4 ratio of recruited T cells extracted from immunoactive ("Full") and control scaffolds. **(C,D)** Flow cytometry analysis of endogenous T cell activation is studied 17 days after subcutaneous implantation of scaffolds. Activation of recruited CD8+ T cells was monitored by measuring surface expression of CD44 as well as intracellular measurement of Granzyme B (GZMB) expression. **(C)** Percentage of T cells with high expression of CD44 and mean fluorescence intensity (MFI) of T cells upregulating CD44 were plotted alongside with representative flow cytometry graphs. **(D)** Percentage of T cells with high intracellular expression of GZMB and MFI of GZMB secreting T cells were plotted. Representative flow cytometry graphs also presented. Each point represents one mouse. Immunoactive scaffold (n = 7) compared with control scaffolds (n = 4).

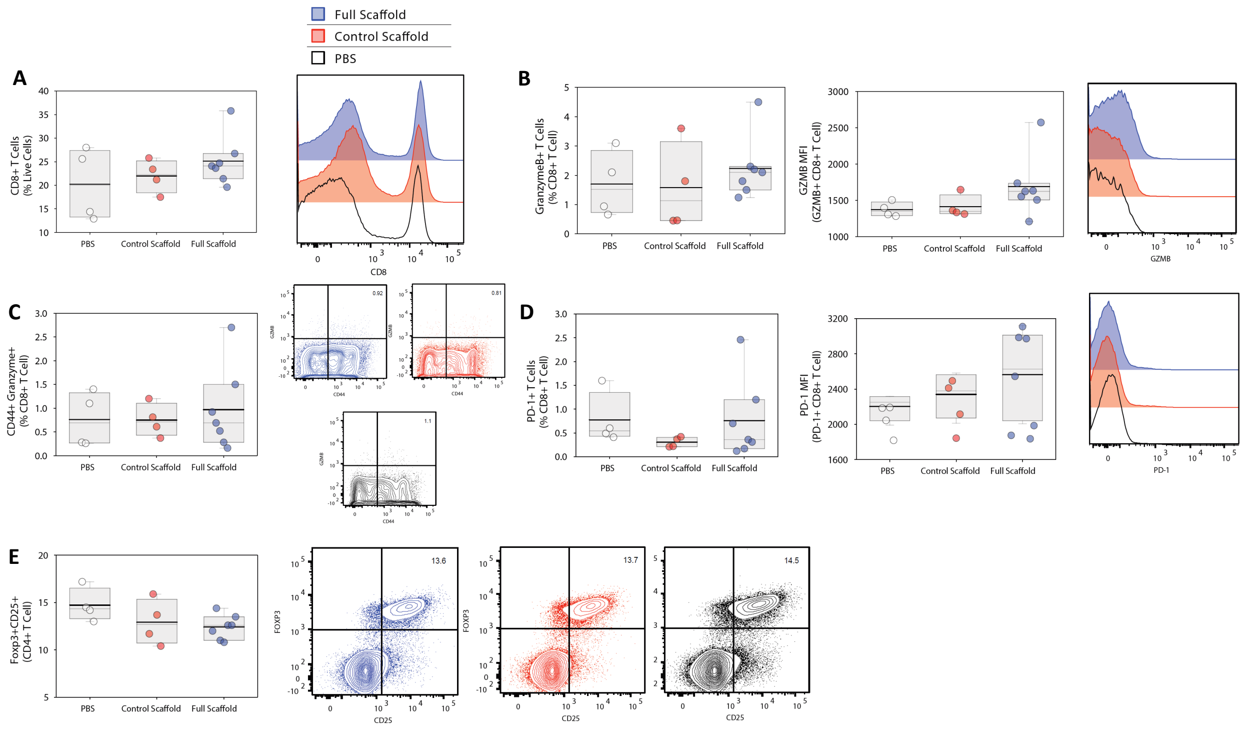

**Extended Data Fig. 7 | No major effects on T cells in tumor-draining lymph nodes. (A)** Flow cytometry analysis of percentage of CD8+ T cells in tumor draining lymph nodes 22 days after subcutaneous injection of B16F10-ova cells in mice receiving different treatment. **(B-D)** Activation of CD8+ T cells in the tumor draining lymph nodes was monitored by measuring their surface CD44 expression as well as Granzyme B (GZMB) intracellular expression. **(B)** Percentage of T cells with high intracellular expression of GZMB and mean fluorescence intensity (MFI) of T cells upregulating GZMB were plotted alongside with representative flow cytometry graphs. **(C)** Percentage of T cells with high expression of CD44 activation marker and ZMB effector cytokine were plotted. Representative flow cytometry graphs also presented. **(D)** Percentage of PD-1 expressing T cells and their MFIs gated on PD-1+ T cells were plotted. Representative flow cytometry graphs also presented. **(E)** The frequency of Foxp3+CD25+CD4+ Tregs in tumor draining lymph nodes. Representative flow cytometry graphs are shown for mice treated with Immunoactive Scaffolds (Blue), Control Scaffolds (Red), and PBS (Black). Each point represents one mouse.

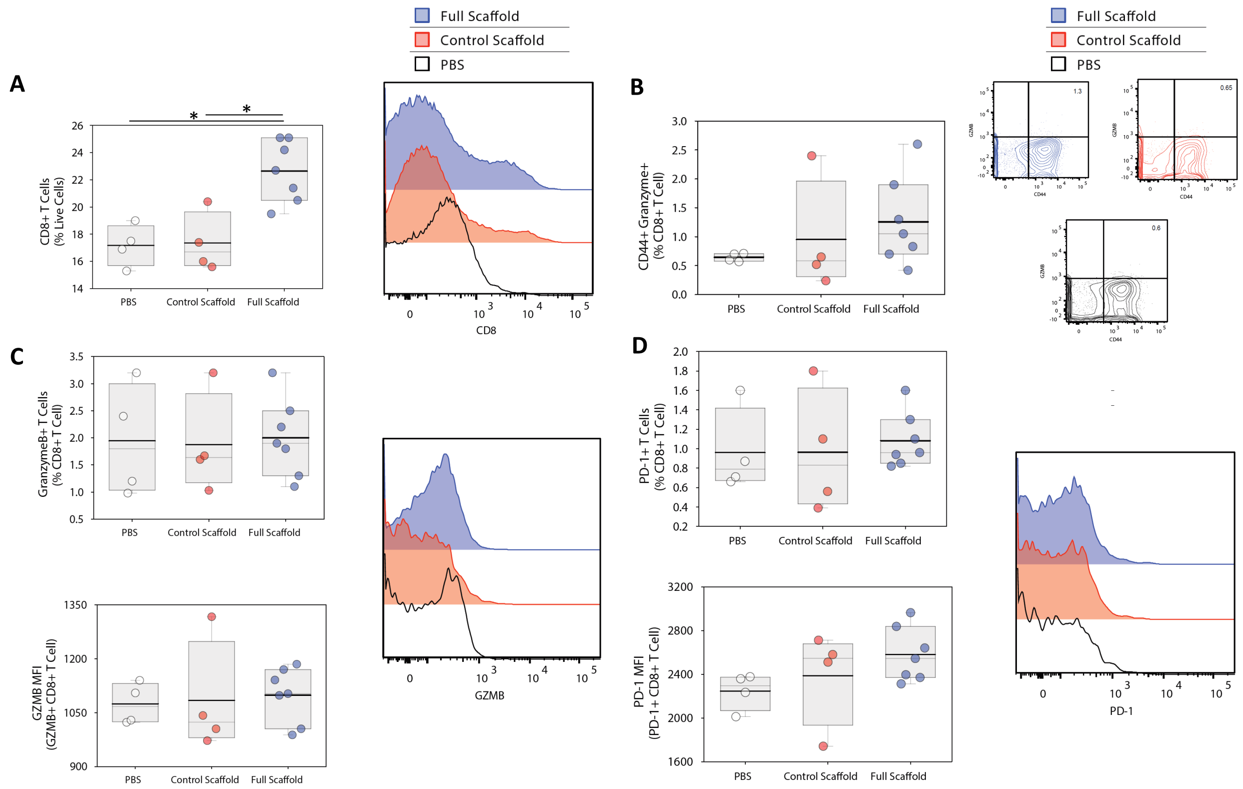


**Extended Data Fig. 8 | No major effects on T cells in CD8+ T cells in Spleen. (A)** Flow cytometry analysis of the percentage of CD8+ T cells in the spleen of tumor-bearing mice 22 days after subcutaneous injection of B16F10-ova cells for mice with different treatments. **(B)** Flow cytometry analysis of T cell activation is studied 22 days after inoculation of tumor cells. Percentage of GZMB+CD44+ T cells was similar in treated vs untreated conditions accompanied with their FACS representatives **(C)** Percentage of T cells with high intracellular expression of GZMB and mean fluorescence intensity (MFI) of T cells upregulating GZMB were plotted alongside with representative flow cytometry graphs. **(D)** Percentage of PD-1 expressing T cells and their MFIs gated on PD-1+ T cells were plotted. Representative flow cytometry graphs also presented. Each point represents one mouse.

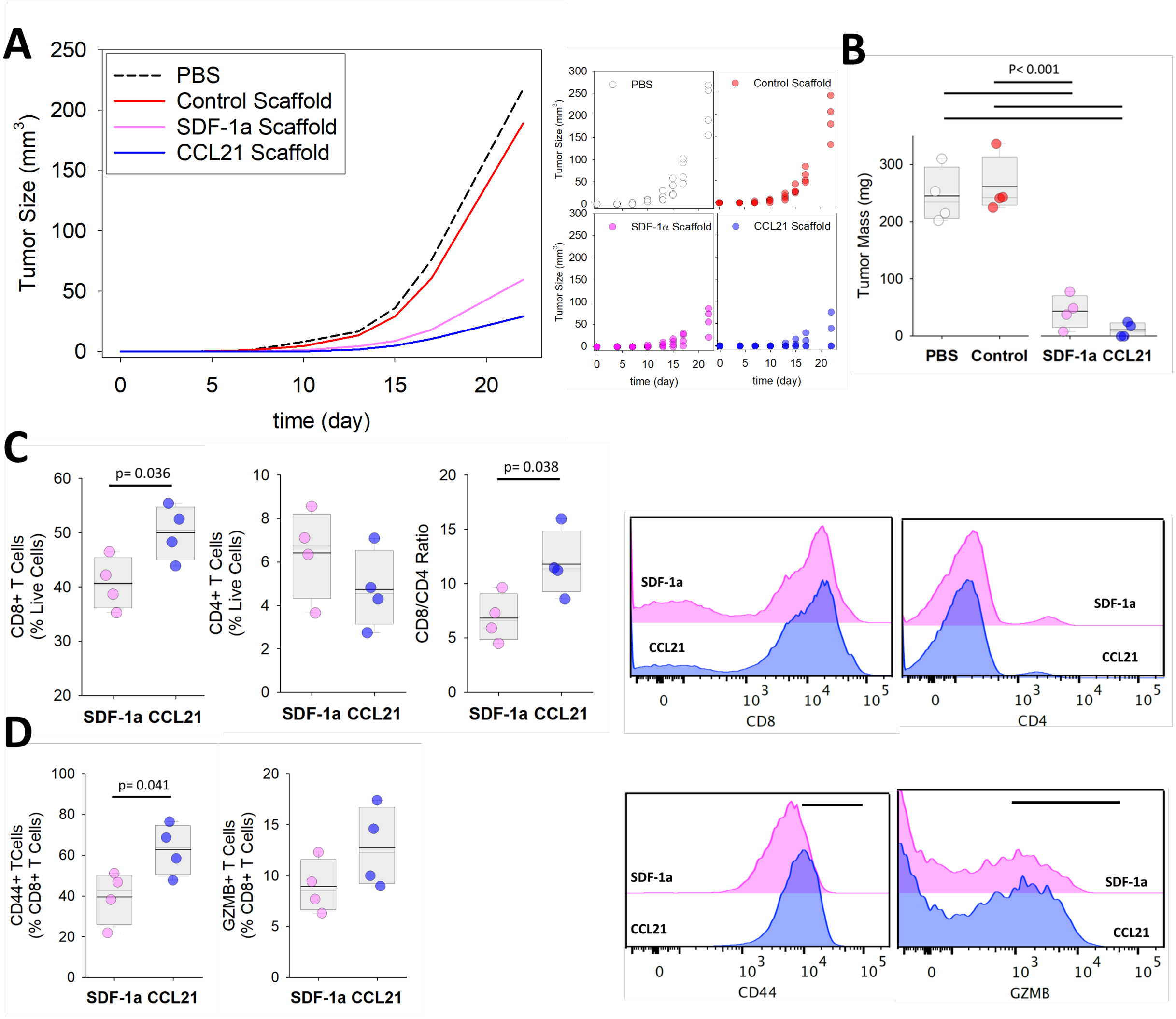


**Extended Data Fig. 9 | Different chemokines in the scaffold can alter the recruitment of endogenous T cells and clearance of tumors. (A)** Melanoma (B16-F10-Ova) tumor growth in wild-type mice with functionalized scaffolds, control scaffolds or PBS treatment (n= 4-7). The therapeutic effects of two chemokines (CCL21 and SDF-1a) were studied by assessing tumor growth over time. **(B)** Tumor masses were measured 22 days after tumor inoculation in wild-type mice treated with functionalized or control scaffolds or PBS treatment. **(C)** Flow cytometry analysis of CD4+ and CD8+ T cells recruited by the scaffolds 17 days after subcutaneous implantation of functionalized scaffolds releasing either CCL21 or SDF-1a chemokines (n=4). FACS quantification of CD8-to-CD4 ratio of recruited T cells extracted from immunoactive and control scaffolds. **(D)** The frequency of activated CD44+CD8+ and GZMB+CD8+ in scaffolds after treating mice with functionalized Scaffolds releasing either CCL21 or SDF-1a chemokines (n=4). Representative flow cytometry data were provided. Each point represents one mouse.

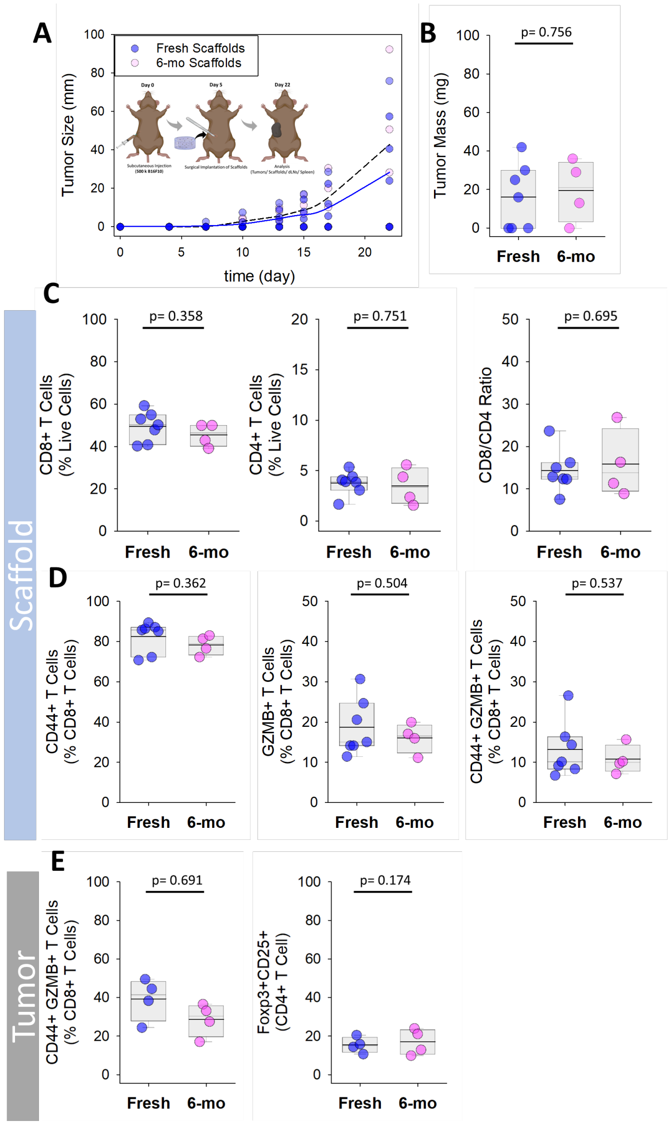


**Extended Data Fig. 10 | Engineered scaffolds can preserve their therapeutic function several months after fabrication. (A)** Melanoma (B16-F10-Ova) tumor growth. Inset: Timing of tumor inoculation and follow up surgical implantation of cells-free scaffolds. **(B)** Final tumor masses were measured 22 days after tumor inoculation in wild-type mice treated with fresh or 6-months old (Immunoactive) scaffolds (n= 4-7). Each point represents a mouse. **(C)** Recruitment and activation of endogenous CD8+ and CD4+ T cells in freshly prepared and 6-Months old scaffolds. Flow cytometry analysis of percentage of CD8+, and CD4+, ratio of CD8+/CD4+ T cells **in the scaffolds**. (D) The frequency of activated CD8+ T cells in the scaffolds assessed by CD44, GZMB, as well as co-expression of CD44 and GZMB T cells in freshly prepared or 6-months old Immunoactive **Scaffolds**. **(E)** The frequency of activated CD44+GZMB+CD8+ and Foxp3+CD25+CD4+ Tregs in **tumors** after being treated with fresh or 6-months old Immunoactive Scaffolds. Each point represents one mouse.

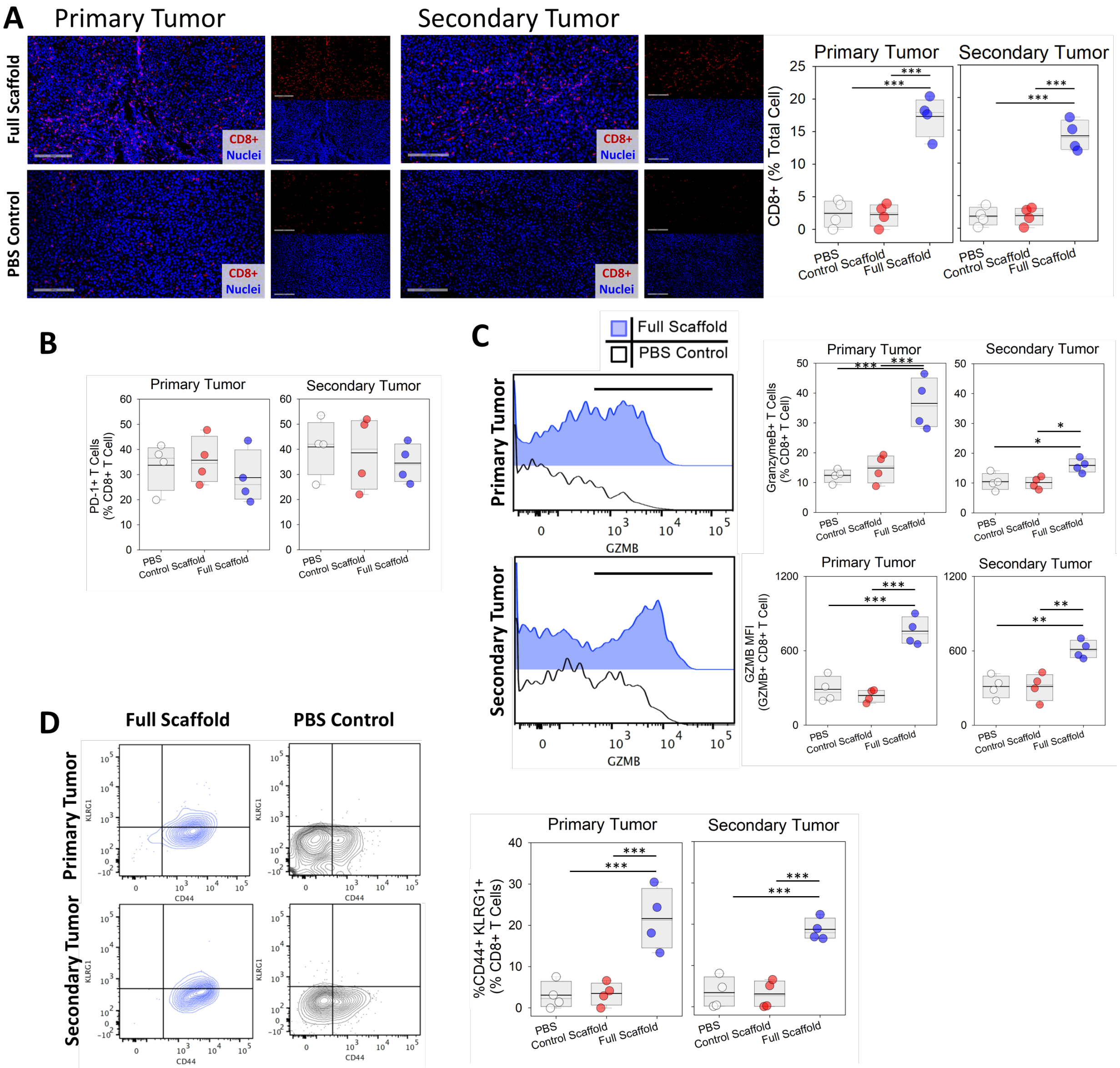


**Extended Data Fig. 11 | Secondary tumor model, T cell activation in the tumors. (A)** Tumor-associated CD8+ T cells were stained in primary and secondary tumors 22 days after tumor inoculation. The frequency of CD8+ T cells **in tumors** were quantified and presented. **(B)** Flow cytometry study of PD-1+CD8+ T cells present in primary and secondary tumors after being treated with immunoactive or control scaffolds. **(C)** Flow cytometry study of GZMB+CD8+ T cell presence in primary and secondary tumors after being treated with immunoactive or control Scaffolds. The frequency and MFI of GZMB+CD8+ T cells in primary and secondary tumors. **(D)** Flow cytometry study of CD44+KLRG-1+CD8+ T cell presence in primary and secondary tumors after being treated with immunoactive or control Scaffolds. Representative FACS and frequency of CD44+KLRG-1+CD8+ T cells in primary and secondary tumors. Note: As 3 out of 7 mice treated with immunoactive scaffold formulation did not grow tumors, these representative sections were only found in the few mice with remaining tumors. (n=4). Each point represents one mouse.

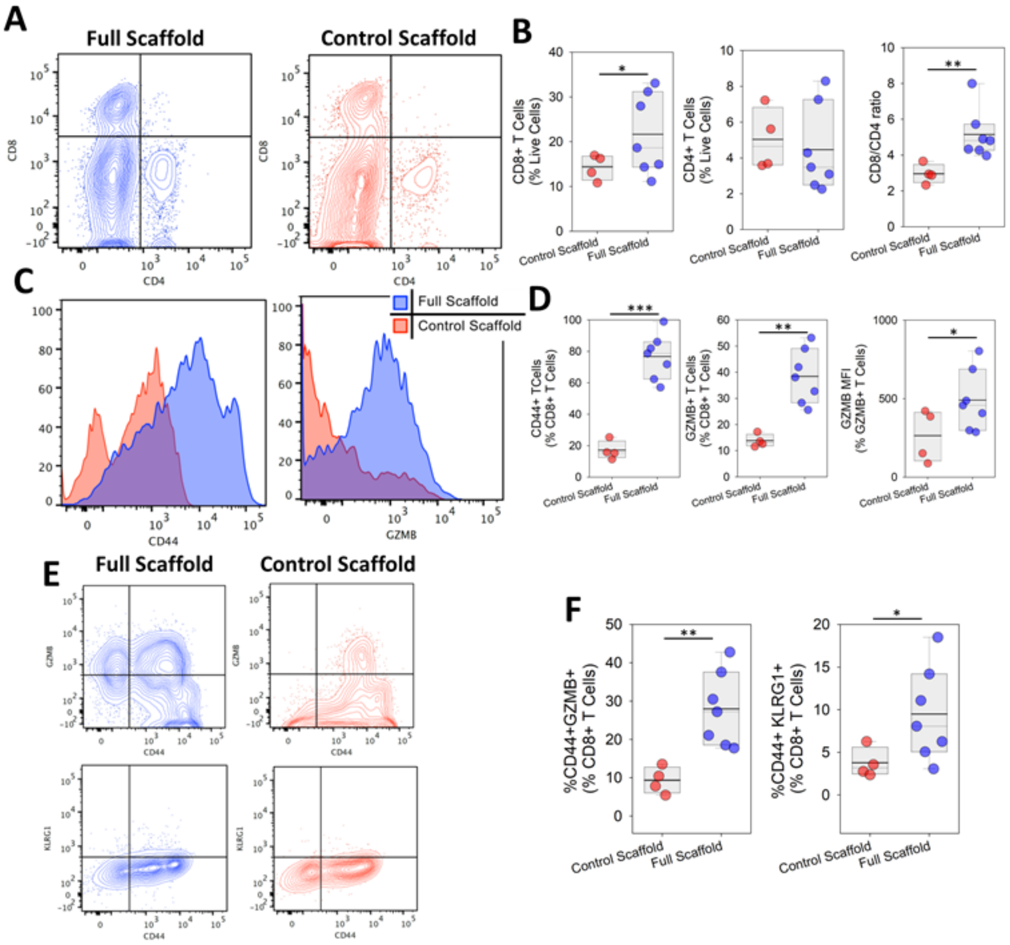


**Extended Data Fig. 12. Secondary tumor model, T cell activation <u>in the scaffolds</u>. (A)** Flow cytometry study of CD8+ T cell presence **in scaffolds** implanted adjacent to tumors. **(B)** The frequency of CD8+ and CD4+ T cells as well as the CD8-to-CD4 T-cell ratios in immunoactive (n=7) and control scaffolds (n=4). **(C-D)** Flow cytometry study of CD44+CD8+ and GZMB+CD8+ T cell presence in scaffolds 17 days after being implanted in tumor-bearing mice. (C) Representative FACS and (D) frequency of CD44+CD8+ and GZMB+CD8+T cells as well as MFI of GZMB+CD8+ T cells in immunoactive (n=7) and control (n=4) scaffolds. **(E-F).** Flow cytometry study of CD44+KLRG-1+CD8+ T cell presence in scaffolds 17 days after being implanted in tumor-bearing mice. (E) representative FACS and (B) the frequency of (F) CD44+KLRG-1+CD8+ T cells in Immunoactive (n=7) and Control (n=4) scaffolds.

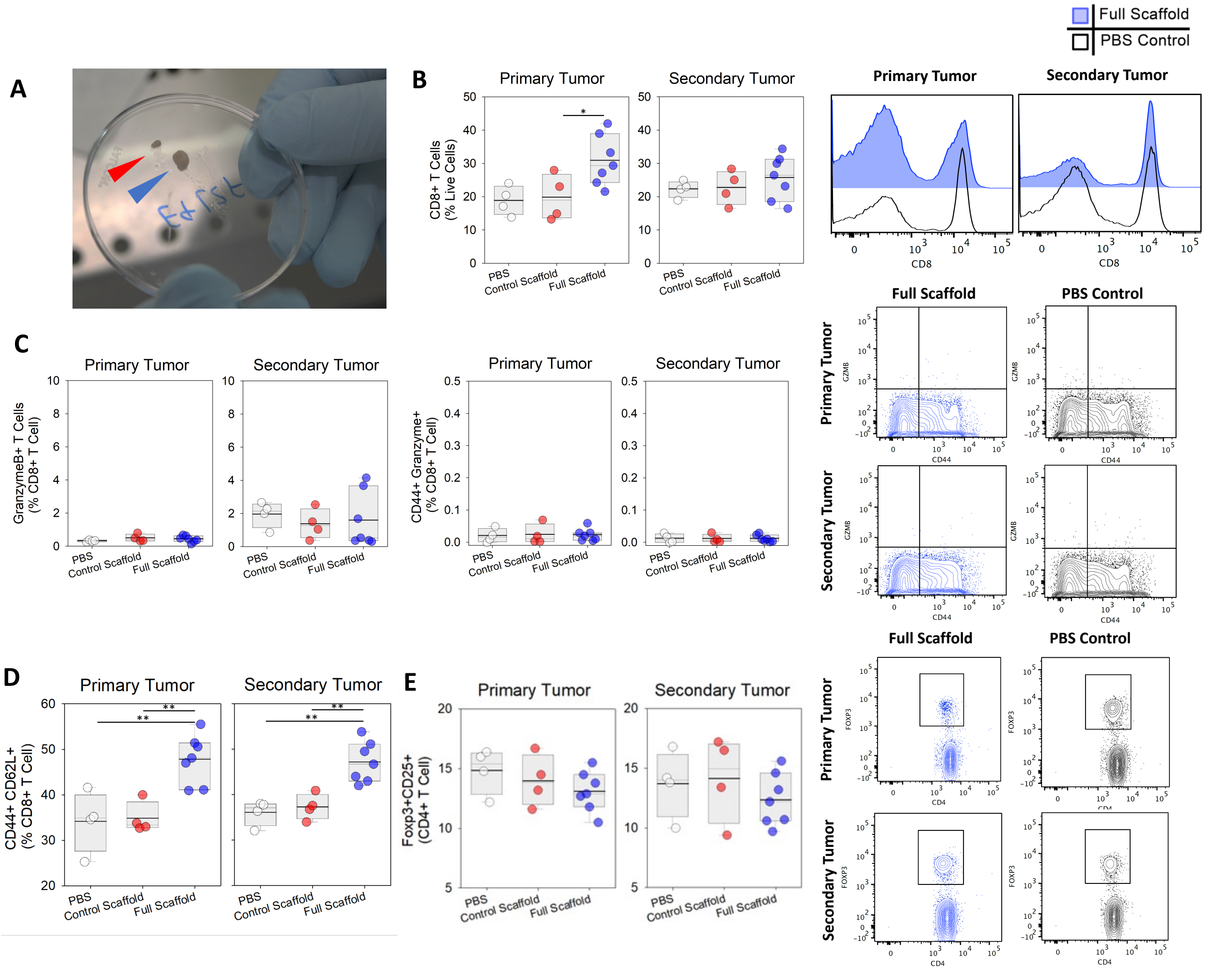


**Extended Data Fig. 13 | Secondary tumor model, T cell activation <u>in the draining lymph nodes</u>**, after being treated with immunoactive (n=7) or control (n=4) scaffolds. **(A)** Showing typical images of primary draining lymph nodes of animal treated with either with immunoactive (blue arrow) or control (red arrow) scaffolds. **(B)** Representative FACS graphs and the frequency of CD8+ T cells in draining lymph nodes of primary and secondary tumors. **(C)** Flow cytometry study of GZMB+CD8+ T cell presence in draining lymph nodes of primary and secondary tumors after being treated with immunoactive (n=7) or control (n=4) Scaffolds. The frequency of GZMB+CD8+ T cells in draining lymph nodes of primary and secondary tumors. Flow cytometry study of CD44+GZMB+CD8+ (effector) T cell presence in draining lymph nodes of primary and secondary tumors after being treated with immunoactive (n=7) or control (n=4) Scaffolds. **(D)** Flow cytometry study of the frequency of CD44+CD62L+CD8+ (central memory) T cell presence in draining lymph nodes of primary and secondary tumors after being treated with immunoactive (n=7) or control (n=4) scaffolds. **(E)** The quantified frequency of Foxp3+CD25+CD4+ Tregs found in tumor draining lymph nodes. Representative flow cytometry of Foxp3+CD25+CD4+ Tregs in primary and secondary tumor draining lymph nodes for mice treated with immunoactive scaffolds (Blue) and PBS (Black).

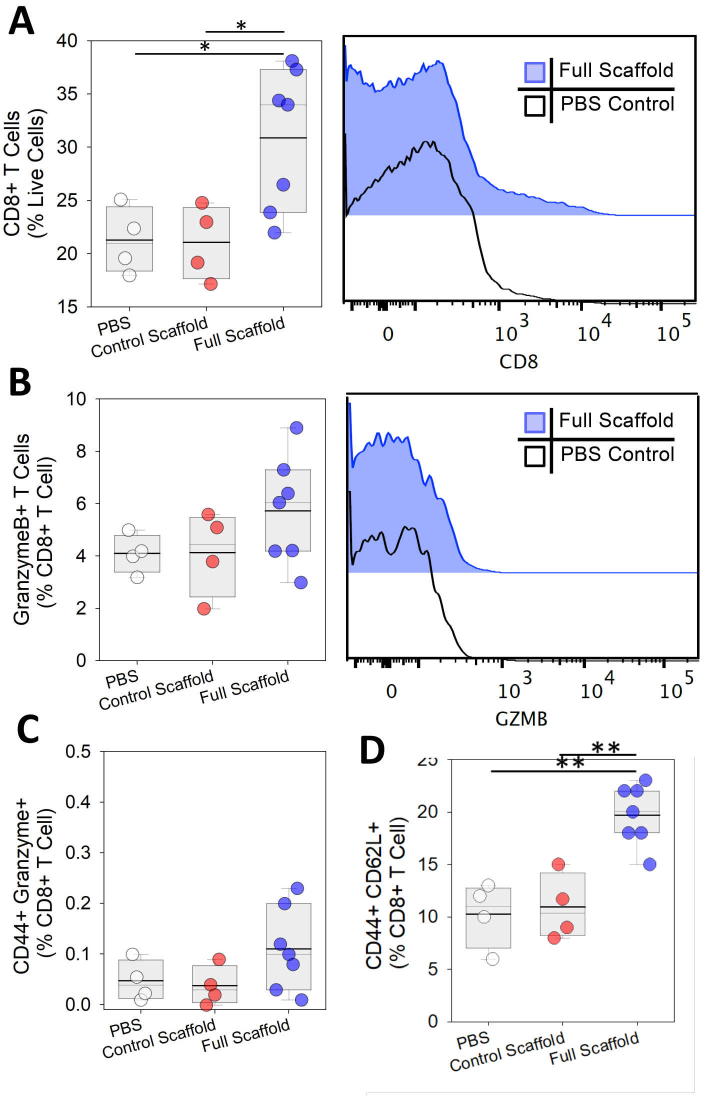


**Extended Data Fig. 14 | Secondary tumor model, T cell activation <u>in the spleen</u>**. Flow cytometry study of CD8+ T cell presence in the spleen of mice after being treated with immunoactive (n=7) or control (n=4) scaffolds. **(A)** The frequency of CD8+ T cells and representative FACS graphs. **(B)** Representative FACS graphs and frequency of GZMB+CD8+ T cells in the spleen. The frequency of **(C)** CD44+GZMB+CD8+ (effector) and **(D)** CD44+CD62L+CD8+ (central memory) T cells in the spleen.

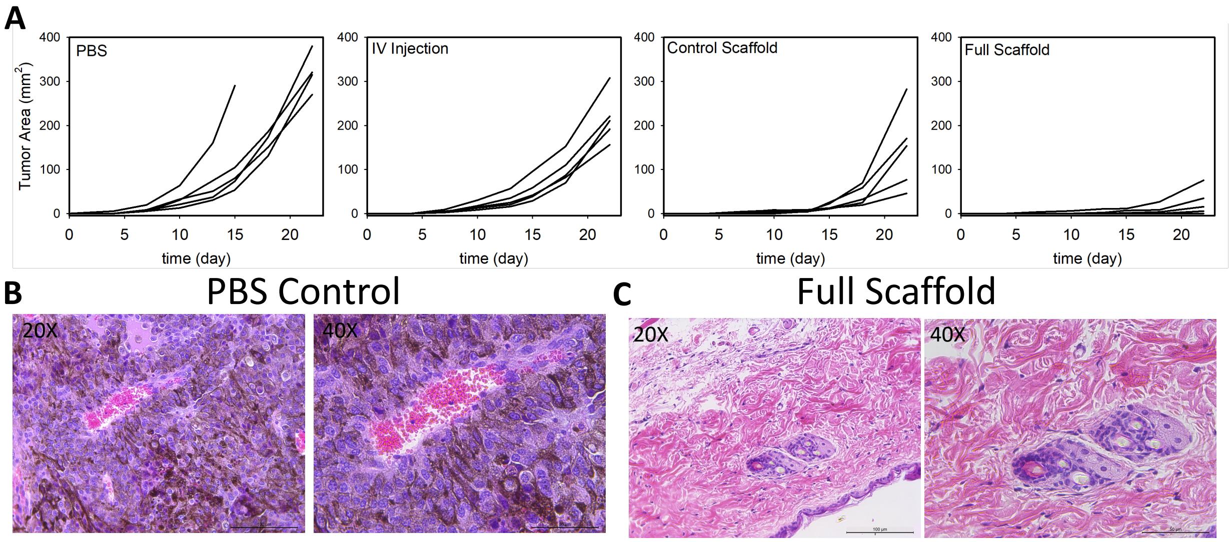


**Extended Data Fig. 15 | Adoptive T cell model, tumor growth and histology. (A)** Melanoma (B16-F10-Ova) tumor growth for groups with different treatments. Each line represents the tumor size of a single mouse over time. **(B-C)** Histologic analysis of the tumor tissues via H&E stain for animals used as **(B)** PBS control *vs.* **(C)** OT-I-loaded Immunoactive Scaffolds.

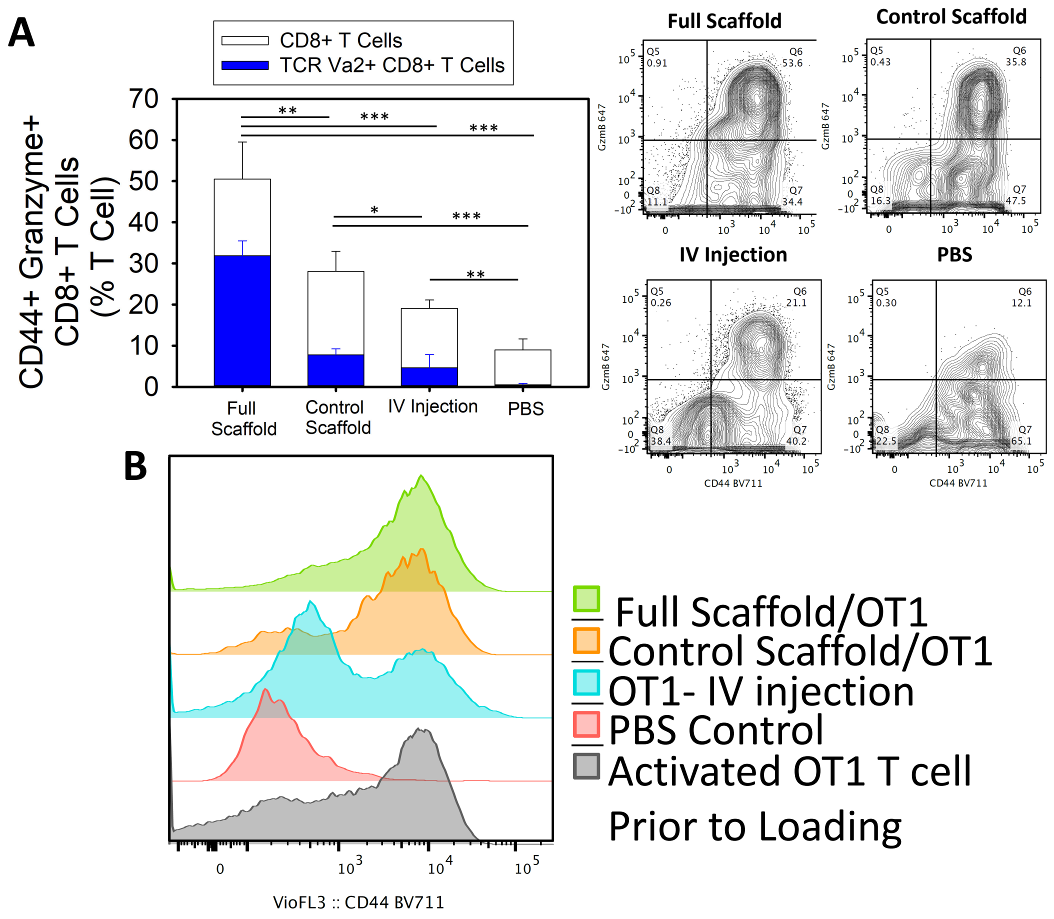


**Extended Data Fig. 16 | Adoptive T cell model, presence of activated CD8+ T cells <u>in tumors</u>.** **(A)** Presence of activated tumor-antigen specific (OT1) CD8+ T cells as well as CD8+ T cells in the tumors was studied 22 days after inoculation of tumor cells using flow cytometry. Percentage of T cells with high expression of CD44 activation marker and GZMB effector cytokine were plotted. Representative flow cytometry graphs also presented. (n= 5). **(B)** Flow cytometry used to identify the presence of activated (CD44 expressing) CD8+ T cells in tumors.

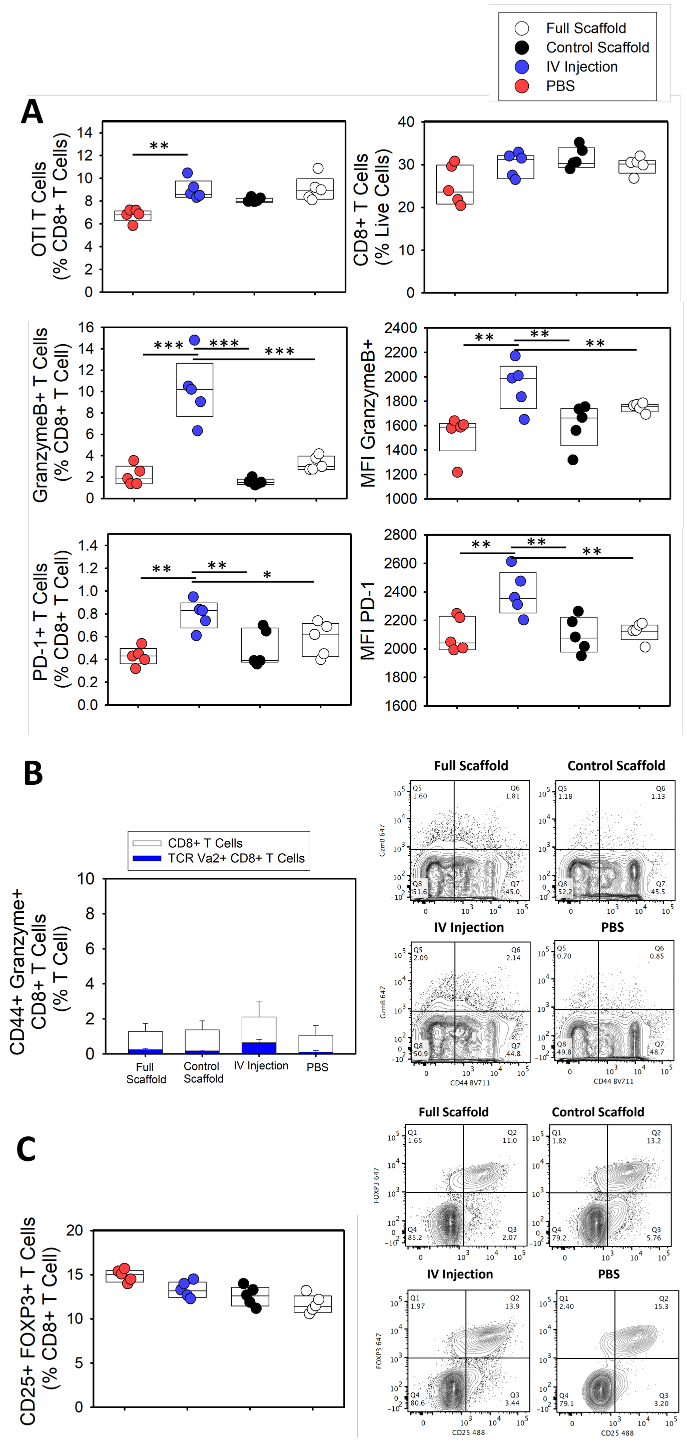


**Extended Data Fig. 17 | Adoptive T cell model, presence of activated CD8+ T cells in <u>tumor draining lymph nodes</u>. (A)** Presence of tumor-antigen specific CD8+ T cells (OTI) as well as activation of CD8+ T cells in the tumor draining lymph nodes was studied 22 days after inoculation of tumor cells using flow cytometry. Percentage of OTI and CD8+ T cells found in tumor draining lymph nodes. Frequency of CD8+ T cells with high expression of GZMB and PD-1 and mean fluorescence intensity (MFI) of T cells upregulating these two proteins were measured. (n= 5). **(B)** Presence of tumor-antigen specific CD8+ T cells as well as activation of CD8+ T cells in the tumor draining lymph nodes was studied 22 days after inoculation of tumor cells using flow cytometry. Percentage of T cells with high expression of CD44 activation marker and GZMB effector cytokine were plotted. Representative flow cytometry graphs also presented. (n= 5). **(C)** The frequency of Foxp3+CD25+CD4+ Tregs in tumor draining lymph nodes were studied. Representative flow cytometry graphs also presented (n= 5).

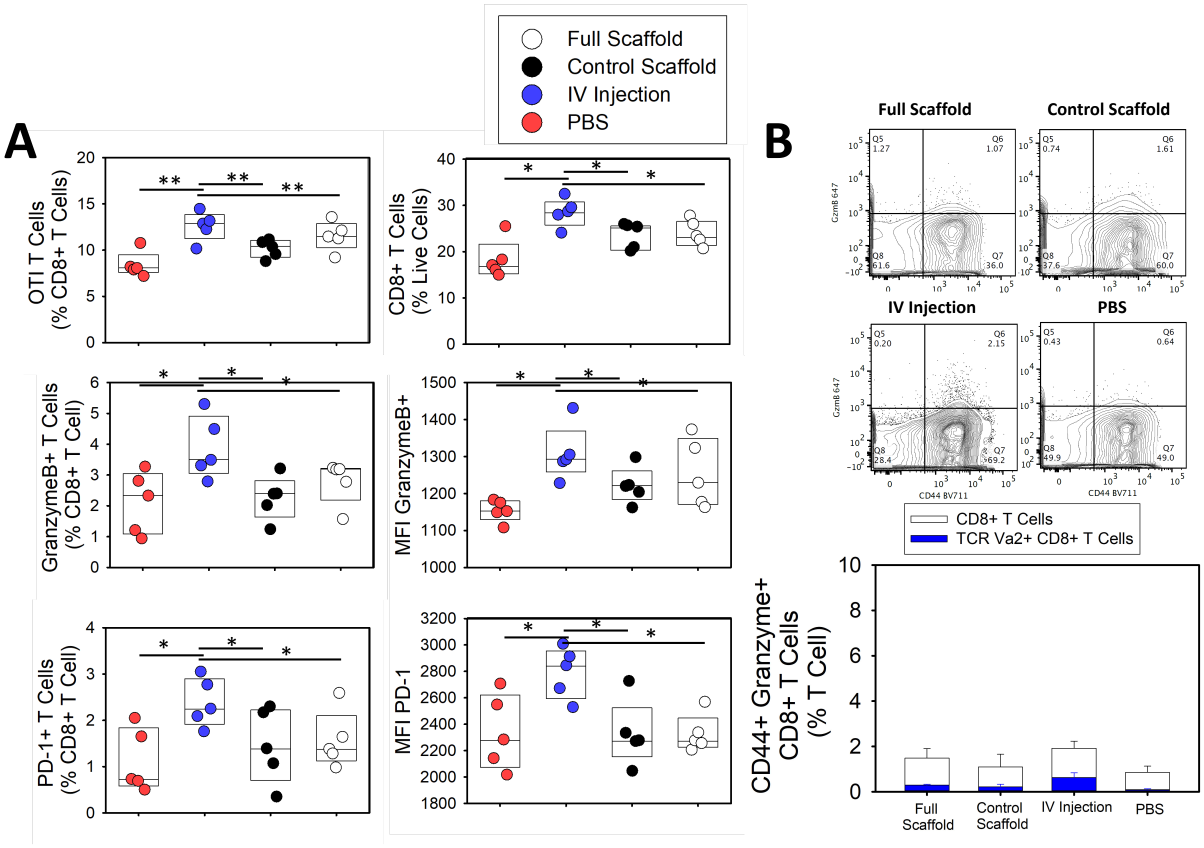


**Extended Data Fig. 18 | Adoptive T cell model, presence of activated CD8+ T cells <u>in spleen</u>. (A)** Presence of tumor-antigen specific CD8+ T cells (OTI) as well as activation of CD8+ T cells in the spleen of tumor bearing mice was studied 22 days after inoculation of tumor cells using flow cytometry. (A) Percentage of OTI and CD8+ T cells found in spleen. Frequency of CD8+ T cells with high expression of GZMB and PD-1 and mean fluorescence intensity (MFI) of T cells upregulating these two proteins were measured. **(B)** Presence of tumor-antigen specific CD8+ T cells as well as activation of CD8+ T cells in the spleen of tumor bearing mice was studied 22 days after inoculation of tumor cells using flow cytometry. Percentage of T cells with high expression of CD44 activation marker and GZMB effector cytokine were plotted. Representative flow cytometry graphs also presented. (n= 5).
